# Activity-regulated micro-exon splicing programs underlie late-onset plasticity at the axon initial segment

**DOI:** 10.1101/2023.10.29.564567

**Authors:** Mohamed Darwish, Satoko Suzuki, Yuki Ogawa, Akinori Takase, Masami Tanaka, Yoko Iijima, Yuji Sato, Mariko Suzuki, Yumi Kanegae, Chisa Okada, Masayuki Takana, Hirotaka-James Okano, Hiroshi Kuba, Takatoshi Iijima

## Abstract

The axon initial segment (AIS) is a specialized neuronal compartment located at the proximal end of axons and initiates action potentials. AIS undergoes plastic changes with aging, disease, and activity levels; however, the molecular mechanisms underlying their plasticity remain unclear. We discovered that depolarization induces diffuse elongation of the AIS in cerebellar granule cells over the span of days via the Ca^2+^-dependent ERK/MAP kinase pathway. These structural changes were accompanied by a decrease in voltage-gated Na^+^ channel density, resulting in a homeostatic attenuation in neuronal excitability. Notably, we found that the late-onset AIS plasticity is associated with depolarization-induced alternative splicing of smaller exons (<100 nt) of transcripts encoding AIS-enriched proteins. In addition, depolarization-induced the skipping of the 53-nt exon19 from the transcript of the splicing protein Rbfox1. CRISPR-mediated removal of exon 19 from Rbfox1 promoted its nuclear localization and sequentially induced a series of downstream micro-exon splicing changes in several AIS proteins, recapitulating cerebellar AIS plasticity. In a Rbfox1-independent mechanism, depolarization-induced insertion of the developmentally regulated micro-exon 34 into the key AIS scaffolding protein Ankyrin G (AnkG). The constitutive insertion of exon 34 into AnkG disrupted its interaction with the AIS cytoskeletal protein βIV spectrin and induced plastic changes in the AIS. Our findings provide fundamental mechanistic insights into the activity-mediated late-onset plasticity of AIS, highlighting the power of micro-scale splicing events in the homeostatic regulation of axonal remodeling.

## Introduction

The axon initial segment (AIS) is a specialized neuronal compartment located at the proximal end of axons and initiates action potentials (1). The function of AIS relies on the local enrichment of a macromolecular complex including several voltage-gated ion channels and cell adhesion molecules (CAMs). These components are assembled by a scaffolding protein, ankyrin-G (AnkG) which associates with cytoskeletal protein, beta-IV (βIV) spectrin, forming a specialized submembranous scaffold at the AIS (2, 3). In particular, the clustering of voltage-gated Na^+^ (Na_v_) channels by AnkG on the AIS is essential for action-potential initiation. Thus, AnkG is critical for AIS assembly and considered as the master organizer of the AIS (4, 5).

The AIS undergoes dynamic changes in structure and function in a homeostatic manner, in response to developmental, pathological, or neuronal activity (6–9). The structural properties of AIS plasticity in the central nervous system (CNS) have been mainly characterized by alterations in AIS length or location (10–15). Notably, in the avian cochlear nucleus, deprivation of auditory input increases AIS length in the nucleus magnocellularis (NM) neurons over several days (10). This elongation enhances neuronal excitability by incorporating Na^+^ conductance to the distal location and, compensating for the loss of auditory nerve activity. In contrast, depolarizing the NM neurons with high K^+^ treatment results in AIS shortening and reduced excitability (16). In hippocampal neurons, chronic depolarization via photo-stimulation or high K^+^ treatment induces a distal shift of the AIS away from the soma resulting in decreased neuronal excitability (11). Thus, AIS plasticity is dynamically regulated bidirectionally and the specific structural changes are highly variable between the neuronal cell types. Despite these findings, the regulatory mechanisms underlying homeostatic AIS plasticity and its function in the mammalian CNS are not fully elucidated.

Several AIS-enriched proteins have multiple alternatively spliced isoforms that are regulated during neural development (17–20), emphasizing the temporal significance of these isoforms in the formation and maturation of AIS. Notably, the Rbfox family (Rbfox1/2/3) orchestrates the alternative splicing (AS) of cytoskeletal and membrane genes encoding AIS proteins and is essential for AIS assembly in ES-derived neurons (18). Disrupting these factors in mice leads to abnormal excitability, resulting in seizures and ataxia (21, 22). These studies highlighted the significance of neuronal alternative splicing programs in AIS formation. Intriguingly, neuronal AS is also modulated via calcium (Ca^2+^)-dependent signaling pathways (23) and controlled by specific RNA-binding proteins in the CNS (19, 24). We recently showed that the Rbfox1 regulates neuronal activity-dependent AS of neurofascin, a neuronal CAM at the AIS, in cerebellar granule cells (CGCs) (19), raising the possibility of its potential impact in modulation of AIS structure and functions through activity-dependent AS (25). The findings have unveiled a promising path toward understanding the mechanisms of neuronal activity-regulated AS and its implications on AIS functionality.

In this study, we demonstrated that activity-regulated AS events contribute to a novel type of homeostatic AIS plasticity in CGCs. Unlike structural AIS plasticity reported previously, prolonged depolarization-induced a diffuse elongation of AIS over several days, ultimately reducing neuronal excitability in a homeostatic manner. Notably, we revealed that neuronal activity-dependent AS of micro/small-exons underlies cerebellar AIS plasticity in both Rbfox-dependent and independent mechanisms. Using genome-editing tools, we demonstrated the causality between micro-exon splicing and AIS structural plasticity in cerebellar neurons.

## Results

### Depolarization induces late-onset structural changes at the AIS in cerebellar granule cells (CGCs)

We initially assessed AIS development in CGCs, a major type of neurons in the cerebellar cortex, and visualized the entire AIS structure by immunostaining against AnkG in cultured neurons after 3 days *in vitro* (DIV3), DIV7 and DIV10. We detected Immunoreactivity at proximal axons in developing CGCs on DIV3 and observed a robust intensity in the highly differentiated cells on DIV7 and DIV10 (data not shown). We then explored whether neuronal activity causes a plasticity-dependent change in differentiated CGCs. We depolarized CGC cultures with 25 mM K^+^ (final concentration, 30 mM) for 3 days on DIV9-DIV12 (Figure 1A and S1A). We found that while AnkG immunoreactivity was robustly concentrated at the proximal position in the untreated CGCs, it was diffusely elongated (approximately three-fold) toward terminal axons in high K^+^ treated CGCs (Figures 1B-1D). Consistently, high-resolution microscopy images with airy scanning showed that the intensity of AnkG staining was robustly reduced at the proximal axons and diffusely elongated with high K^+^ stimulation (Figures 1C and 1D). We detected similar structural changes in the AIS when we checked the distribution of other AIS proteins such as Neurofascin and βIV spectrin upon K^+^-induced depolarization (Figure S1A and S1B) or when neuronal activity was increased in a more physiological way by bicuculine, a competitive antagonist of GABA A receptors (Figure S1C). Together, these data revealed a novel form of activity-dependent structural change at the cerebellar AIS.

**Figure 1.**
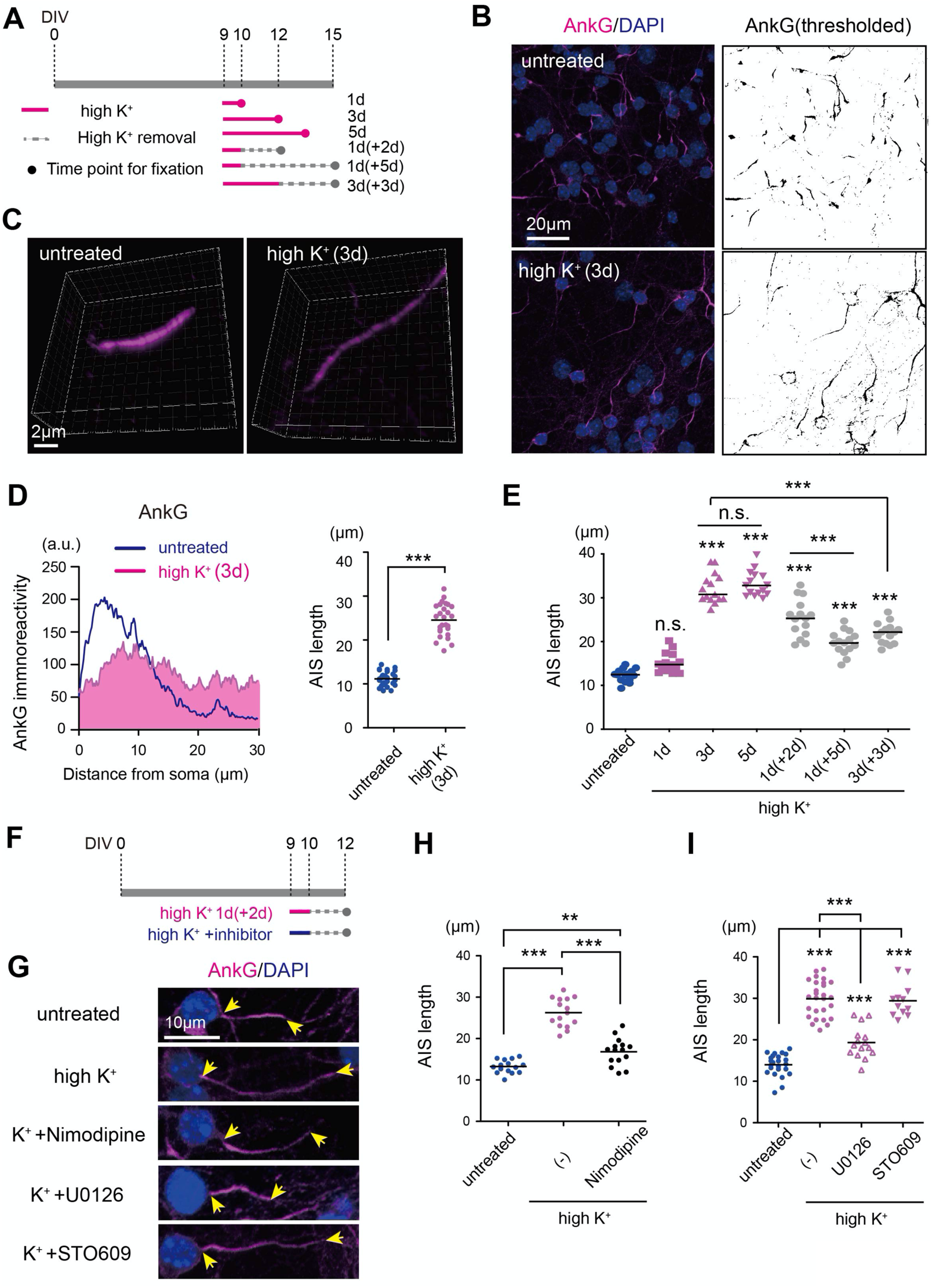
Depolarization-induced robust change in AIS structure of CGCs. (A) Schematic representation showing the time course of high K+ stimulation in CGCs. The cultures were maintained in normal K+ media (5mM, control; gray bar) and high K+ media (final concentration, 30mM, intervention; red bar) from days in culture (DIV) 9 for 1 day or 3 days, with or without recovery 2-5 days after K+ stimulation. (B) Immunostaining of AnkG in untreated and high K+-stimulated CGCs for 3 days (3d) as described in A. DAPI stains the nucleus. (C) Representative high magnification Airyscan images of AnkG in untreated and high K+-treated CGCs (3d). (D) Left, Representative fluorescence intensity of AnkG staining in untreated and high K+-treated CGCs described in B. The intensities of more than 10 axons were averaged per group. Right, Quantification of AIS length of neurons in B. [n = 30 fields (control) and 27 fields (high K+-treated)], unpaired t-test. (E) Quantification of AIS length of CGCs treated with high K+ stimulation at different durations and conditions as described in A. (n = >15 fields for each group). One-way ANOVA with Tukey’s multiple comparisons test. (F) Schematic representation showing the time course of the experiment. The CGCs were simultaneously treated with high K+ and inhibitors for 1 day and maintained for an additional 2 days in a normal medium [1d+(2d) as shown in Figure 1A]. (G) Representative image of CGC AIS, untreated, or treated with high K+ stimulation alone or with each of the following inhibitors: Nimodipine, STO-609 (10 µM), and U0126 (5 µM). The arrowheads mark the start and the end of the AIS. (H) Quantification of AIS length (n = 15 fields for each group). (I) Quantification of AIS length (n = 24, 27, 15, and 12 fields for [control, high K+, K++U0126, and K++STO609, respectively). The scale bars represent 20 μm in (B), 2 μm in (C), and 10 μm in (G).

To assess the time scale and characteristics of depolarization-induced structural changes in the AIS, we exposed CGC cultures to high K^+^ stimulation for different durations (1 day, 3 and 5 days) and conditions (Figure 1A). We found that AIS elongation was nearly saturable with high K^+^ stimulation for 3 days and no structural changes were observed after 1 day. Although the 1-day treatment did not induce elongation in AIS, maintaining the culture for an additional 2 days in a normal medium was sufficient to induce robust AIS elongation comparable to that of the CGC neurons stimulated for 3 days (Figure 1E) suggesting that transient stimulation within at most 1 day would be sufficient for exerting a maximum change in the AIS structure. In addition, when CGCs were further maintained in a normal medium for more than 3 days after removal of medium with high K^+^ (i.e., 1 day or 3 days), the AIS length was significantly restored (Figure 1E). Together, these data indicate that the depolarization-induced changes in the AIS structure of CGCs are late-onset, long-lasting, and reversible. We then tested the contribution of Ca^2+^ influx to the AIS structural changes. The simultaneous application of nimodipine, an inhibitor of the L-type voltage-dependent Ca^2+^ channel (L-VDCC) strongly abolished the elongation of AIS induced by high K^+^ treatment (Figure 1F-G). We then assessed the signaling pathways downstream of Ca^2+^ that underlie neuronal activity-mediated AIS structural changes. We performed pharmacological inhibition of calcium/calmodulin-dependent protein kinase (CaMK) and extracellular signal-regulated kinase/mitogen-activated protein kinase (ERK/MAPK) simultaneously with high K^+^ treatment. Remarkably, AIS changes were significantly blocked by ERK/MAPK, inhibitor U0126, while there were no significant changes in response to CaMK kinase inhibitor, STO609 (Figure 1F and 1H). Thus, these data reveal that extracellular Ca^2+^ influx via L-VDCC and subsequently activated downstream ERK/MAPK signaling triggers long-term structural change at the AIS in CGCs.

### Depolarization-induced structural changes in CGC AIS are accompanied by reduced density of Na^+^ channels and attenuated neuronal excitability

We next investigated whether the activity-regulated structural changes at the AIS in matured CGCs are accompanied by functional and physiological changes. Given that neuronal activity causes the AIS to undergo a dynamic change in structure and function (10, 11), we tested whether applying high K^+^ stimulation could cause some plasticity-dependent changes at the AIS in CGCs. The distribution and density of voltage-gated Na^+^ channels (Na_v_) regulate the generation of action potential. Since AnkG is necessary for the clustering of Na_v_ at the AIS (26), we hypothesized that AnkG structural changes in the K*^+^*-treated neurons could alter the clustering of Na_v_ and hence neuronal excitability. Immunostaining with a pan-Na_v_ antibody that recognizes all Na^+^ channel subtypes, revealed that Na_v_ was distributed diffusely along the axons in high K^+^-treated cultures, whereas it was densely concentrated on the proximal part in the control cultures (Figure 2A and 2B). In addition, the high-resolution microscopy imaging with Airyscan exhibited that Na_v_ was highly co-localized with AnkG in the axons even after high K^+^ treatment (Figure 2C), indicating that the altered distribution of Na_v_ is largely followed by that of AnkG. We next investigated the biophysical consequences of activity-regulated structural changes at the AIS using electrophysiological studies.

**Figure 2.**
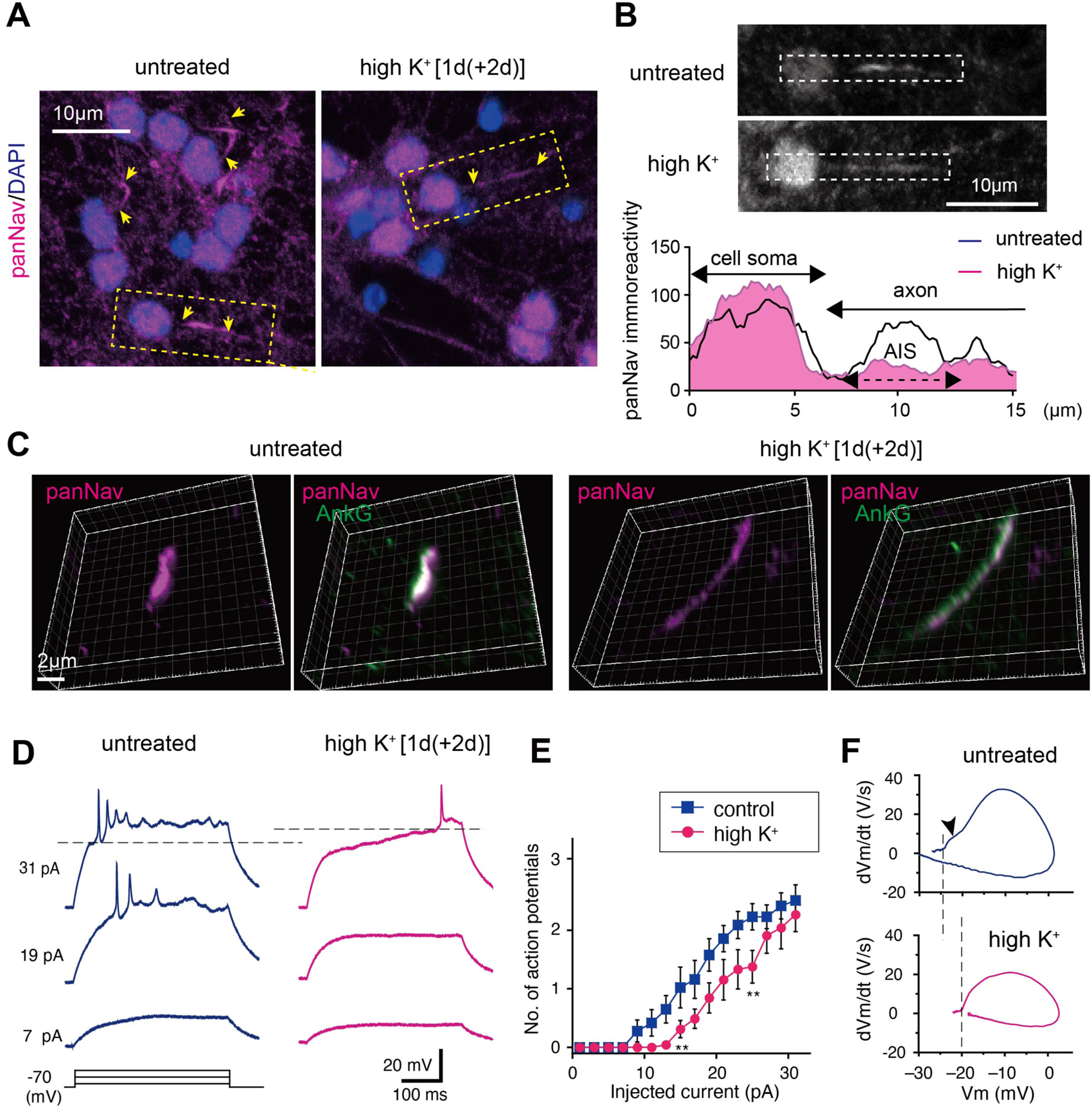
Depolarization-induced structural changes in the AIS reduce the density of the Na^+^ channel at the AIS and dampens neuronal excitability. (A) Immunostaining of pan-Nav in untreated and high K+-stimulated CGCs for 3 days. DAPI stains the nucleus. (B) Upper: High magnification images of the area surrounded by dashed rectangles in A. Lower: Representative fluorescence intensity of pan-Nav staining in untreated and high K+-treated CGCs described in A. (C) Representative high magnification Airyscan images of the co-localization of pan-Nav and AnkG in untreated and high K+-treated CGCs. (D) Voltage responses to current pulses. Depolarizing currents were injected into the soma while maintaining the membrane potential at –70 mV. Traces at 7, 19, and 31 pA are shown in the bottom, middle, and top, respectively. The threshold voltage is indicated by horizontal broken lines. The CGCs are treated with high K+ [1d+(2d) in Figure 1A]. (E) The relationship between the number of action potentials and the injected current for the control cells (n = 22 and 24 cells for the control and high K+-treated group, respectively). (F) Phase plot showing the dVm/dt and Vm relationship of the action potential in (E). The threshold voltage is indicated by vertical dashed lines. The upstroke of the action potential is biphasic for untreated CGCs and monophasic for high K+-treated CGCs. The scale bars represent 10 μm in (A and B), 2 μm in (C), and 10 μm in (G).

We injected a depolarizing current into the soma of CGCs and recorded the action potentials under a current clamp. The current threshold was elevated in the high K^+^-treated cultures (16.3±1.0 pA and 21.9±1.0 pA in untreated (n=22) and treated CGCs (n=24), respectively (p=0.0008) (Figure 2D). Therefore, the number of action potentials decreased in the treated cultures, which led to a rightward shift of the input-output curve (Figure 2E). The maximum dV/dt of the action potentials was decreased in the treated cultures, which indicated a decreased Na^+^ current: the action potentials were 26.3±2.0 V/s and 18.7±1.7 V/s for untreated (n=22) and treated CGCs (n=24), respectively (p=0.006). Moreover, the phase plot of the action potential upstroke was biphasic in the untreated CGCs, but the initial phase was absent in the treated CGCs (arrowhead in Figure 2F). suggesting that the axonal component of the action potential was suppressed after the high K^+^ treatment. Thus, the diffusion of Na_v_ on the axon could contribute to the attenuation of neuronal excitability in high K^+^-treated cultures. We also checked the total amount of Na_v_ proteins in high K-treated CGCs and the levels were comparable to untreated groups (Figure S1D). Together these findings suggest that depolarization-induced structural and functional changes occur at CGCs AIS contribute to controlling the neuronal activity in a homeostatic fashion.

### Depolarization dynamically alters AS of small/micro-exons of transcripts encoding AIS-enriched proteins in CGCs

To comprehensively explore the molecular changes occurring during AIS plasticity at the transcriptome-wide level, we conducted RNA-seq analyses on high K-stimulated CGCs and untreated controls. In high K^+^ stimulated neurons, we identified 222 differentially expressed genes (DEGs), of which 137 genes were upregulated and 85 genes were downregulated (threshold set: FC>4.0, corrected p<0.05) (Figure S2A). Gene Ontology (GO) analyses exhibited that the DEGs were highly enriched in terms of calcium and synapse and KEGG pathway enrichment analysis showed that DEGs were enriched to the MAPK signaling pathway (Figure S2B). Actually, we observed a lot of previously reported calcium-sensitive transcripts were significantly altered in the treated neurons: calcium-dependent synaptic organizers *Cbln1*, *Cbln3,* and neuronal pentraxin (*Nptx1*) were dramatically reduced, while brain-derived neurotropic factor (*Bdnf*) was increased (27) (Figure S2C). Although high K^+^ stimulation successfully induced the alteration of neuronal activity-related genes, it didn’t induce significant changes in the transcript of major AIS-related proteins (Figure 3A and S2D).

**Figure 3.**
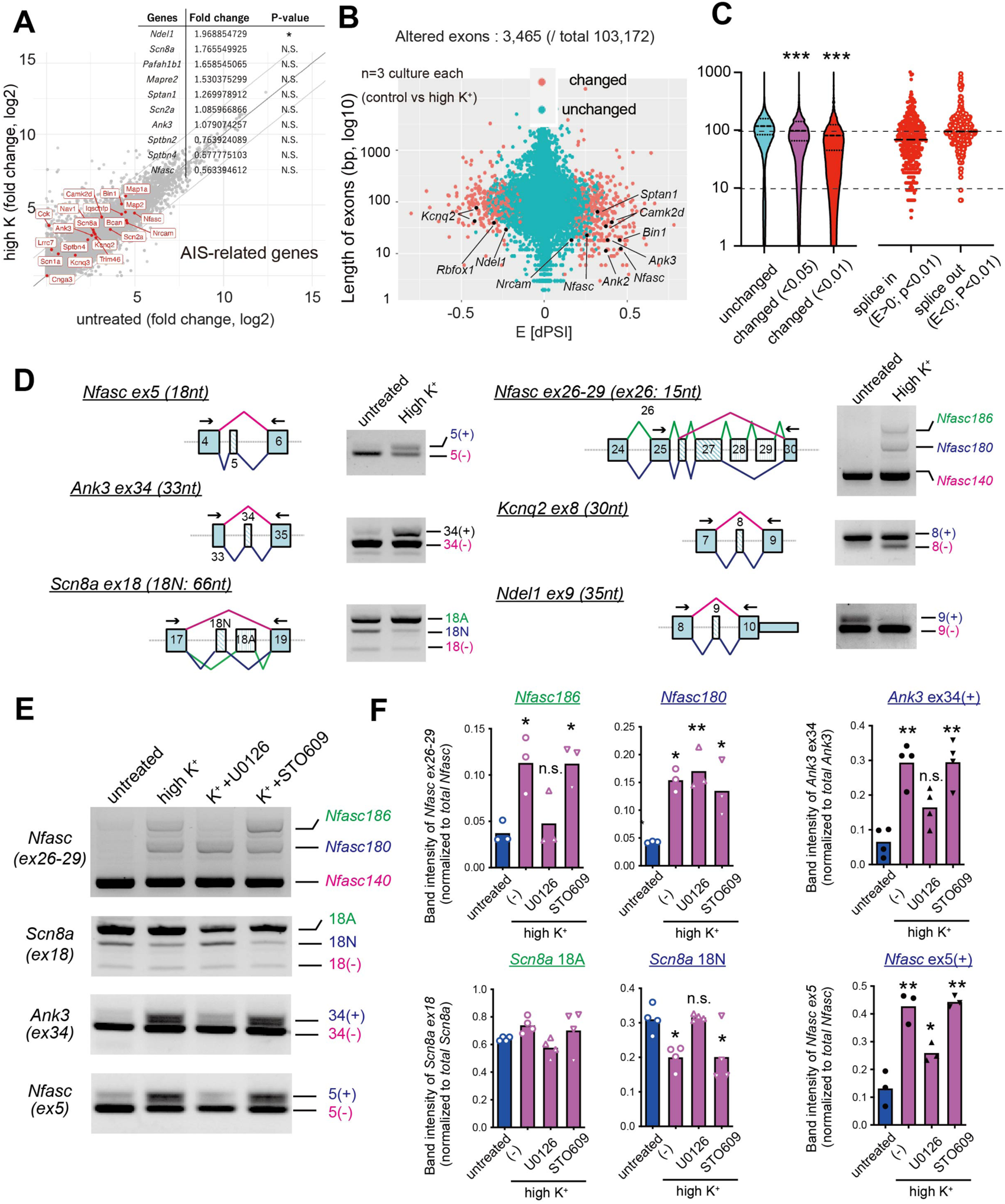
The altered small/micro-exon splicing of AIS-related genes in cultured CGCs upon depolarization. (A) Scatter plot of the RNA-seq data showing no significant differences in the expression levels of genes encoding AIS-related proteins (highlighted in red) between untreated and high K+ (3d) CGCs. The mean fold changes (FC) and p-values of AIS-related genes are presented in the table. n = 3 cultures from three independent experiments. (B) Correlation between the exon length (Y-axis) and splicing alteration (X-axis), represented as delta PSI (dPSI). Red dots indicate significantly altered exons, while black dots represent significantly altered exons of AIS-related genes. (C) Violin plots illustrate the correlation between exon length and splicing alteration in response to high K+ treatment. Smaller exons (<100nt) display greater susceptibility to significant splicing changes, with the spliced-in exons generally being shorter than the spliced-out exons. (D) Semi-quantitative RT-PCR validation of depolarization-induced small/micro-exon splicing changes in genes encoding AIS-enriched proteins. Schematic illustration of AS choice is depicted in the left panels, and representative gel images are presented in the right panels. (E) Semi-quantitative PCR images demonstrating the splicing changes in genes encoding AIS-enriched proteins in CGCs, either untreated or treated with high K+ stimulation alone or with each of the following inhibitors: STO-609 (10 µM) and U0126 (5 µM). (F) Densiometric quantification of alternatively spliced exons in the groups shown in (E). n = 3-4 independent cultures per group. One-way ANOVA with Tukey’s multiple comparisons test.

We therefore dissected the data with the single exon scale using vast-tools, a computational approach that quantifies AS and identifies differentially spliced AS events using RNA-seq data (28). Among the total 103,172 exons identified, 3,465 exons, including those of AIS proteins, exhibited significant alterations in the treated CGC neurons (Figure 3B). Notably, we observed a trend where smaller exons (<100nt) were more susceptible to significant splicing changes (Figure 3C and S2E), highlighting the correlation between neuronal activity-dependent splicing and exon length. Additionally, the spliced-in exons tended to be shorter than the spliced-out exons (Figure 3C and S2E), emphasizing the influence of neuronal activity on the splicing process and exon size regulation. Interestingly, we found that alternative small/ micro-exons of transcripts encoding AIS-enriched proteins are preferentially altered in the treated CGCs (Figure 3B, black dots). These splicing changes included the insertion of exon 34 (33nt) of the transcript encoding core AIS component *Ank3* (AnkG) and exons 5 (18nt) and 26 (15nt) of the adhesion molecule *Nfasc.* Additionally, the exclusion of exon 18N (66nt) from *Scn8a* that encodes the sodium channel *Nav1.6* was observed (Figures 3D and S3). These results indicate that neuronal activity-mediated cerebellar AIS plasticity is accompanied by small/micro-exons splicing changes of the major AIS-related transcripts.

Since depolarization-induced structural AIS changes in the CGCs are mediated through ERK/MAPK signaling (Figure 1F-G), we tested whether this pathway is involved in the activity-dependent AS of small/micro-exons at the AIS-related transcripts. We found that the MEK inhibitor, U1026, but not the CaMK inhibitor STO609, blocked the splicing shifts in some of the transcripts encoding AIS-related proteins such as *Ank3, Nfasc,* and *Scn8a* (Figure 3E and 3F). These data suggest that ERK/MAPK signaling is responsible for not only activity-mediated cerebellar AIS structural plasticity but also the accompanying splicing changes in transcripts encoding AIS-enriched proteins.

### The nuclear Rbfox1 isoform triggers subsequent AS changes in small/micro-exons at the targeted AIS transcripts and modulates cerebellar AIS plastic changes

We next examined the mechanism underlying neuronal activity-dependent splicing of genes encoding AIS-enriched proteins. A previous study showed that depolarization represses exon 19 of the splicing regulator Rbfox1 prompting its nuclear localization to increase its splicing activity (Figure 4A) (29) and we recently demonstrated that activity-dependent nuclear localization of Rbfox1 enhances the inclusion of *Nfasc* exons 26-29 in CGCs (19). We, therefore, asked whether the nuclear Rbfox1 (exon 19-skipped isoform) regulates activity-dependent AS changes in small/micro-exons at AIS-enriched transcripts and underlies AIS structural plasticity. RNA-seq and PCR analyses confirmed a significant exclusion of the 53 nt exon 19 of *Rbfox1* in high K^+^ stimulated CGC neurons (Figure S4A and S4B).

**Figure 4.**
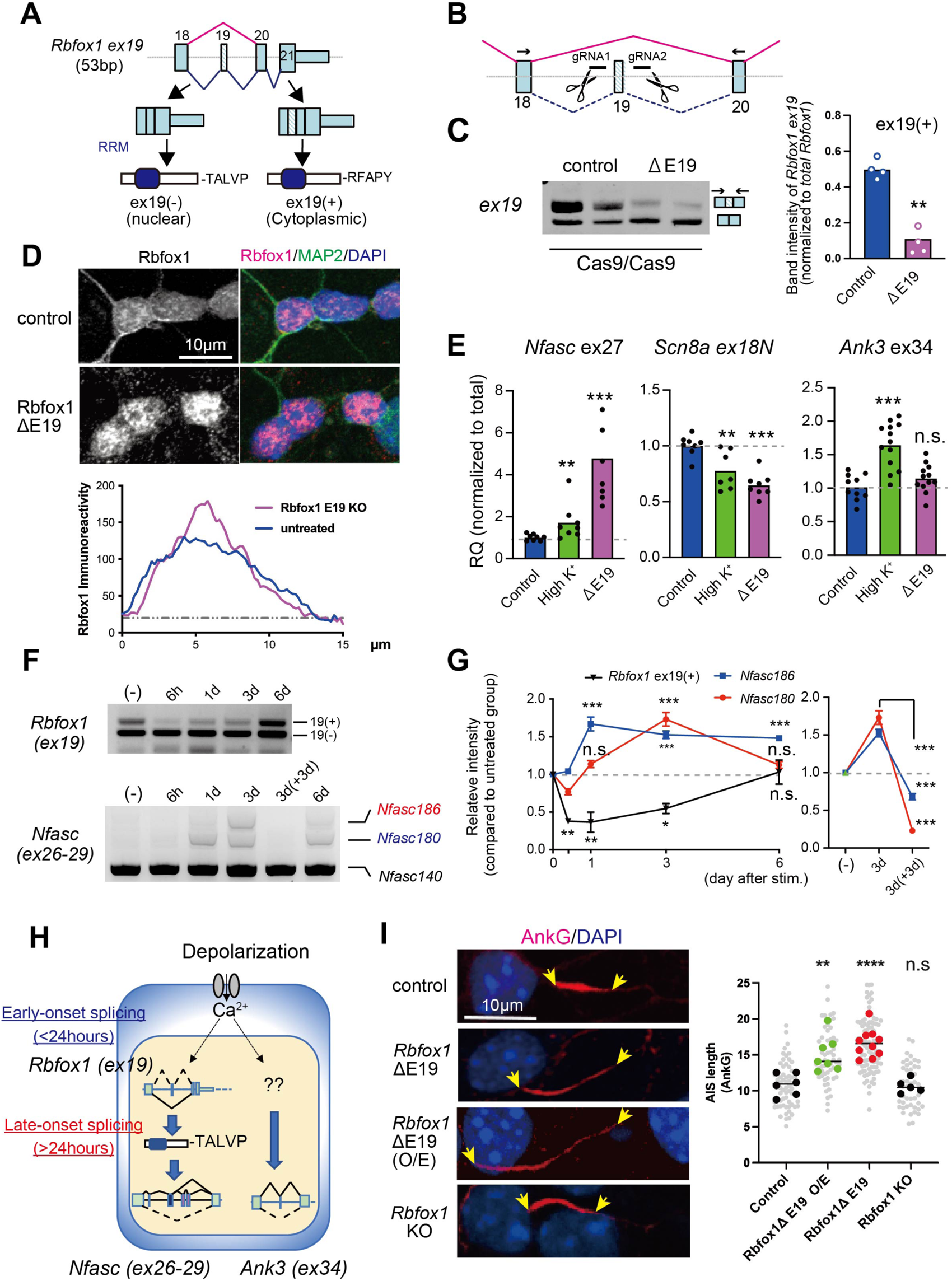
The nuclear isoform of the Rbfox1 splicing factor [Rbfox1 ex19(-)] triggers AS change in downstream target small/micro-exons at AIS transcripts and modulates the late-onset plastic change at the AIS. (A) Schematic representation illustrating the alternative splicing (AS) of exon 19 in Rbfox1 transcripts. The removal of exon 19 alters the reading frame at the C-terminus, resulting in the production of a nuclear isoform of Rbfox1. (B) Schematic diagram demonstrating the removal of *Rbfox1* exon 19 (Δex19) using CRISPR/Cas9-mediated genome engineering. Lentiviral vectors carrying two gRNAs targeting the neighboring intronic sequences of exon 19 are transduced in Cas9-expressing CGCs. (C) Semi-quantitative RT-PCR images (left) and densitometric quantification (right) showing the removal of Rbfox1 exon 19. n = 4 cultures, unpaired t-test. (D) Immunostaining (upper) and representative fluorescence intensity profile (lower) of Rbfox1 in control and Rbfox1 Δex19 CGCs. MAP2 and DAPI stain the entire neuronal morphology and nucleus, respectively. The intensities of more than 10 axons were averaged per group. (E) Relative expression of *Nfasc* exon 26-29, *Scn8a* exon 18 (18N), and Ank3 exon 34 in control and Rbfox1 Δex19 CGCs. The Ct values were normalized to those of the total transcripts. n = >8 cultures per group. One-way ANOVA with Tukey’s multiple comparisons test. (F) Semi-quantitative RT-PCR images showing the splicing changes of Rbfox1 exon 19 and Nfasc exon 26-29 in CGCs treated with high K+ for different durations and conditions as shown in the panel. (G) Densitometric quantification of the spliced exons in the groups shown in (F). The value of the untreated culture was arbitrarily defined as 1.0. n = 4 cultures per group. The data are presented as the mean ± the standard error of the mean (SEM). One-way ANOVA with Dunnett’s multiple comparison test. (H) Model depicting Rbfox1 dependent and independent mechanisms regulating the late-onset AS controls of AIS-related transcripts (I) Representative image of AIS (left) and quantification of AIS length (right) from control, Rbfox1 Δex19, Rbfox1 exon19 (-) O/E, and Rbfox1 knockout CGCs. The arrowheads mark the start and the end of the AIS. n = 6-10 fields per group. One-way ANOVA with Tukey’s multiple comparisons test. The scale bars represent 10 μm in (D and I).

To decipher the role of *Rbfox1* exon 19 in the activity-driven splicing alteration of AIS-related transcripts, we employed CRISPR/Cas9-based-genome editing to remove exon 19 (Figure 4B). We transduced lentiviral vectors carrying gRNAs into CGCs expressing Cas9 and confirmed the successful removal of exon 19 (Figure 4C). Notably, the deletion of exon 19 (△ex19) induced a shift in the subcellular localization of Rbfox1 toward the nucleus. While control CGCs displayed widespread Rbfox1 distribution throughout the cells with immunoreactivity detected on dendritic processes, the Rbfox1 △ ex19 cultures remarkably exhibited condensed immunoreactivity in the nuclei (Figure 4D), simulating the effect of high K^+^ stimulation as reported previously (19, 29).

We next investigated the consequences of Rbfox1 nuclear localization on the splicing program of genes encoding AIS proteins. Strikingly, the △ex19 led to a splicing alteration in various AIS-related transcripts including *Nfasc* exon 26-29 and *Scn8a* exon 18, resembling the effect of high K^+^ stimulation while it does not affect the splicing of *Ank3* exon 34 (Figure 4E). Similarly, the knockdown of Rbfox1 by siRNA abolished the activity-induced shifts in splicing of *Nfasc* exons 26-29 and *Scn8a* exon 18 but did not influence those of *Ank3* exon 34 and *Nfasc* exon 5 (Figure S4C-F). These results highlight the regulatory role of the nuclear *Rbfox1* in the alternative splicing of specific AIS proteins, but it does not fully account for all splicing events during AIS plasticity, suggesting the potential involvement of other splicing regulations in this form of plasticity.

We also examined the temporal splicing changes of *Rbfox1* exon 19 and its targeted transcripts in high-K^+^ treated CGCs under different durations and whether it occurs concomitantly with AIS plasticity. The exon19 was rapidly excluded within less than 6 hours after high K^+^-stimulation and re-inserted up to the initial level for 6 days even upon high K^+^ condition (Figure 4F and 4G), indicating that exclusion of exon19 is early-onset, but transient and reversible. Contrarily, activity-induced shifts in splicing of *Nfasc* exons 26-29, were slower and late-onset, taking several days but the splicing shift was recovered 3 days after high K^+^ removal (Figure 4F and 4G). Strikingly, these splicing events closely mirrored the timing of depolarization-induced structural AIS changes demonstrated in Figure 1E. Together these data suggest that the rapid removal of the activity-regulated exon 19 from *Rbfox1* triggers a delayed splicing shift in targeted transcripts encoding AIS-enriched proteins such as *Nfasc* (Figure 4H) and therefore underlies the late-onset AIS structural plasticity.

Subsequently, we investigated the impact of *Rbfox1* △ex19 on structural AIS plasticity. Interestingly, we found that AnkG staining was reduced at the proximal axon and diffusely elongated the AIS in CGCs lacking *Rbfox1* exon 19 (Figure 4I). Remarkably, the overexpression (O/E) of Rbfox1△ex19 in CGCs also induced similar structural changes in the AIS. In contrast, the knockout of Rbfox1 by removing a constitutive exon failed to prompt significant AIS plasticity alteration (Figure 4I). These data emphasize that *Rbfox1* exon 19 acts as a key driver for multiple splicing events of genes encoding AIS-localized proteins, governing the late-onset AIS plasticity of CGCs.

### Activity-regulated insertion of micro-exon 34 at the neuronal AnkG isoform contributes to depolarization-induced structural changes at the AIS

The AIS structural plasticity driven by nuclear Rbfox1-mediated splicing alterations (Figure 4) was relatively modest compared to the depolarization effect (Figure 1), suggesting that Rbfox1 regulation alone may not fully elucidate the depolarization-induced changes in AIS. We, therefore, focused on “Rbfox1-independent” mechanisms that underlie depolarization-induced AIS plasticity. We showed that depolarization induced the insertion of the micro-exon 34 into AnkG and this splicing shift is independent of Rbfox1 (Figure 4E and S4D and F). We next examined the time course for AnkG exon 34 splicing changes upon depolarization. The Insertion of exon 34 was barely detectable 6 hours after high K^+^ treatment, gradually increasing up to 3–6 days (Figure 5A and 5B) and was completely reversed 3 days after removal of high K^+^ (Figure 5A and 5C). This temporal profile indicated that the activity-dependent splicing events of exon 34 reflected the late-onset and reversible nature of the plastic AIS changes shown in Figure 1E. We, therefore, hypothesized that AnkG exon 34 is a potential player that underlies AIS plasticity. AnkG mainly exists in four spliced isoforms in the CNS: the 190 kDa, 230 kDa, 270 kDa, and 480 kDa forms of AnkG (30–32). The 480 kDa AnkG (AnkG480) is a neuron-specific isoform that is essential for protein assembly at the AIS (33), whereas the 190 kDa AnkG (AnkG190) is a ubiquitously expressed. Exon 34 of AnkG exists at the embryonic stage and is almost completely excluded postnatally (18, 20), encoding a micro-segment of the spectrin-binding domain (SBD) (Figure 5D, red box). To investigate the impact of micro-exon 34 on the localization of AnkG isoforms into AIS, we transduced the neuron culture with viral vectors carrying GFP-fused AnkG480 or AnkG190 with/without exon 34 (Figure 5D). Strikingly, AnkG480 ex34 (-) was properly localized at the proximal AIS position, whereas AnkG480 ex34 (+) exhibited a diffuse distribution toward terminal axons (Figure 5E-5G). In contrast, AnkG190 with or without exon 34 showed comparable axonal localization as both variants were evenly distributed throughout the axons (Figure 5E and 5F). Hence, these findings highlight the physiological implication of activity-regulated exon 34 insertion into neuron-specific AnkG480 in mature neurons.

**Figure 5.**
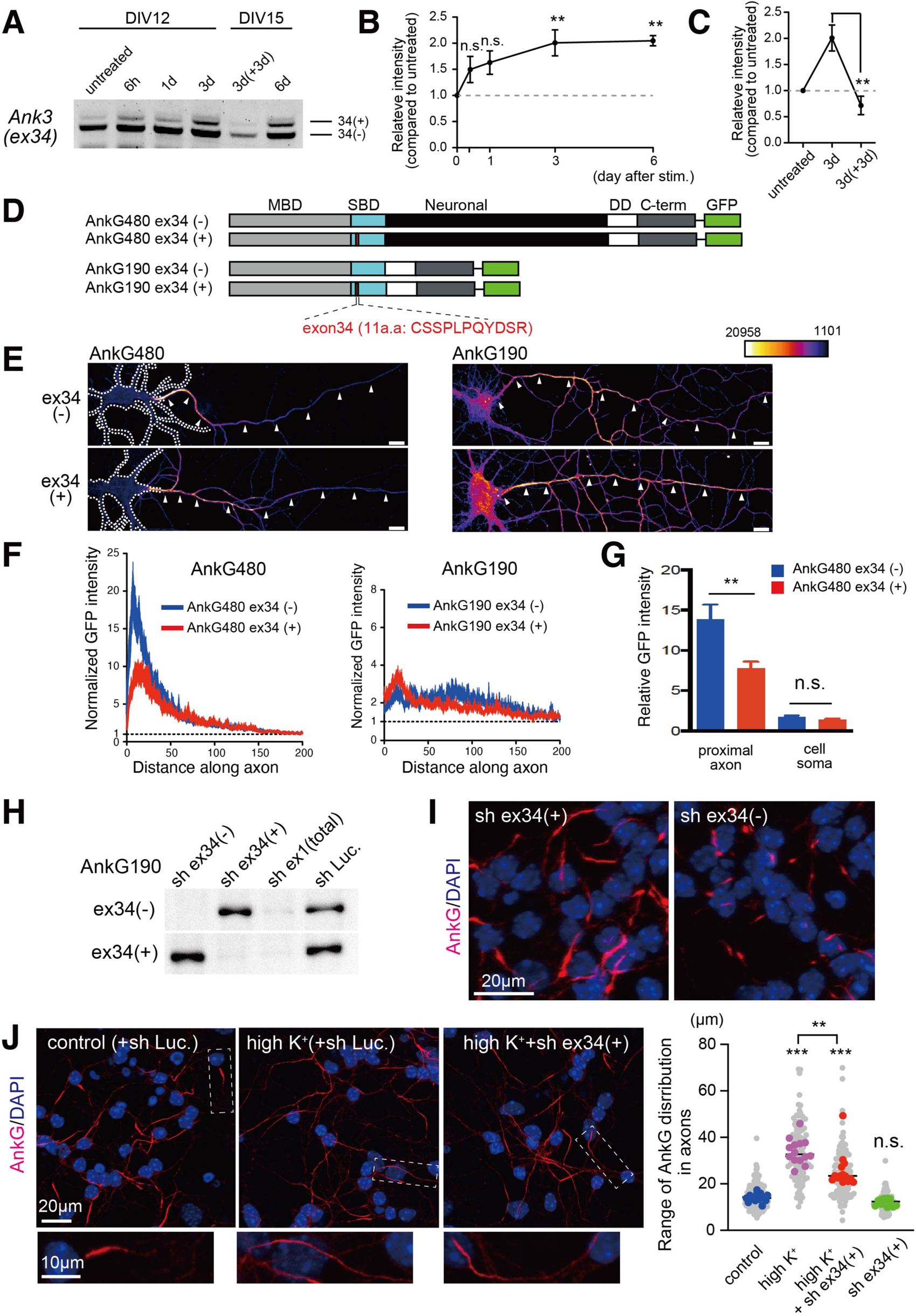
Activity-regulated *Ank3* exon 34 impairs the localization of neuron-specific AnkG at the AIS. (A) Semi-quantitative RT-PCR images showing the splicing changes of *Ank3* exon34 in CGCs treated with high K+ for different durations and conditions as shown in the panel. (B-C) Densitometric quantification of *Ank3* exon34 in the groups shown in (A). n = 3-4 cultures per group. Data are presented as the mean ± SEM. One-way ANOVA with Dunnett’s multiple comparison test. (D) Schematic representation of adenoviral plasmid constructs for AnkG480 and AnkG190 highlighting the position of exon 34 in red boxes. (E) Representative images of cultured hippocampal neurons transduced with indicated adenoviral vectors and immunostained with anti-GFP antibody. (F) Representative fluorescence intensity of GFP immunoreactivity along the axons. (G) Quantification of GFP fluorescent intensity in the proximal axon and soma. n= 11-15 cells per group. Two-way ANOVA. (H) Immunoblot showing AnkG exon 34 (+/-) isoform-specific knockdown with lentiviruses carrying short hairpin RNAs (shRNAs). (I) Immunostaining of AnkG in CGCs transduced with lentiviral vectors carrying shRNAs for AnkG exon 34 (+/-) isoforms. (J) Immunostaining of AnkG in CGCs transduced with lentiviral vectors carrying shRNA for luciferase (control) or AnkG exon 34 (+) with and without high K+ stimulation. High-magnification images of the area surrounded by dashed rectangles are shown. n = 14 fields (untreated), 14 fields (high K^+^-treated), 14 fields (high K^+^-treated + shAnk3 ex34) and 12 fields (untreated + shAnk3 ex34). One-way ANOVA with Dunnet’s test. The scale bars represent 10 μm in (E and J, high magnification), and 20 μm in (I and J, low magnification).

To elucidate the role of activity-regulated insertion of exon 34 in AIS structural changes, we analyzed AIS plasticity in AnkG ex34(+/-) isoform-specific knockdown CGCs. Immunoblot analysis confirmed that the transduction of lentiviral vectors expressing shRNA targeting AnkG with or without ex34 (+/-) [sh ex34(+/-)] dramatically and specifically abolished relevant protein expression in HEK293T cells (Figure 5H). As expected, the disruption of AIS assembly was evident in sh ex34(-), but not sh ex34(+), isoform-specific knockdown CGC neurons (Figure 5I), suggesting that AnkG ex34(-) is the predominant isoform CGCs AIS. We next investigated the effect of AnkG ex34(+) isoform-specific knockdown on high K^+^-treated CGCs to directly examine the effect of activity-regulated insertion of exon 34 on AIS plasticity. Notably, the knockdown of AnkG ex34(+) markedly attenuated the depolarization-induced diffusion of AnkG throughout the axons (Figure 5J). These results suggest the contribution of activity-regulated insertion of AnkG exon 34, in parallel to the sequential splicing events mediated by nuclear Rbfox1, to the structural change during AIS plasticity.

### The constitutive insertion of micro-exon 34 at AnkG gene impairs the interaction of AnkG and βIV spectrin mimics the structural change during cerebellar AIS plasticity

To validate the impact of exon 34 at AnkG on AIS structural changes *in vivo*, we generated AnkG exon 34 knock-in mice (*Ank3* ex34 KI) in which exon 34 is constitutively inserted. We fused exon 34 with the downstream constitutive exon on the *Ank3* genomic locus using the CRISPR/Cas9 system and confirmed that exon 34 is inserted at adult stages (Figure 6A). Exon 34 is a cassette micro-exon that encodes a segment of the spectrin-binding domain (SBD) (Figure 5D) and has been suggested to disrupt AnkG-βIV spectrin physical interaction (18). Co-immunoprecipitation experiment exhibited a robust association between AnkG with βIV spectrin in WT brain, whereas this interaction was severely weakened in *Ank3* ex34 KI brains (Figure 6B), demonstrating the influence of exon 34 insertion on the interaction between AnkG and βIV spectrin. Subsequently, we assessed AIS structure in *Ank3* ex34 KI mice and found that akin to depolarization, AnkG exhibited a diffuse distribution toward terminal axons in the CGCs (Figure 6C). Additionally, the immunoreactivity of the βIV spectrin was reduced but remained largely confined to the proximal portion of *Ank3* exon34 KI neurons (Figure 6D) mirroring the changes observed in high K^+^-induced CGC neurons (Figure S1C) where βIV spectrin immunoreactivity reduced but remained mostly at the proximal portion. Co-immunostaining of βIV spectrin with neurofascin, an AnkG-anchored protein, revealed noticeable disparities in their distributions. While the staining was largely overlapping in the wild-type, a notable mismatch was observed particularly on the distal AIS part with neurofascin exhibiting diffuse distribution along axons in the *Ank3* ex34 KI (Figure S5), consistent with the altered AnkG localization. These data suggest that exon 34 insertion at AnkG not only disrupts its axonal distribution but also its anchoring surface proteins in the AIS, which triggers AIS structural changes.

**Figure 6.**
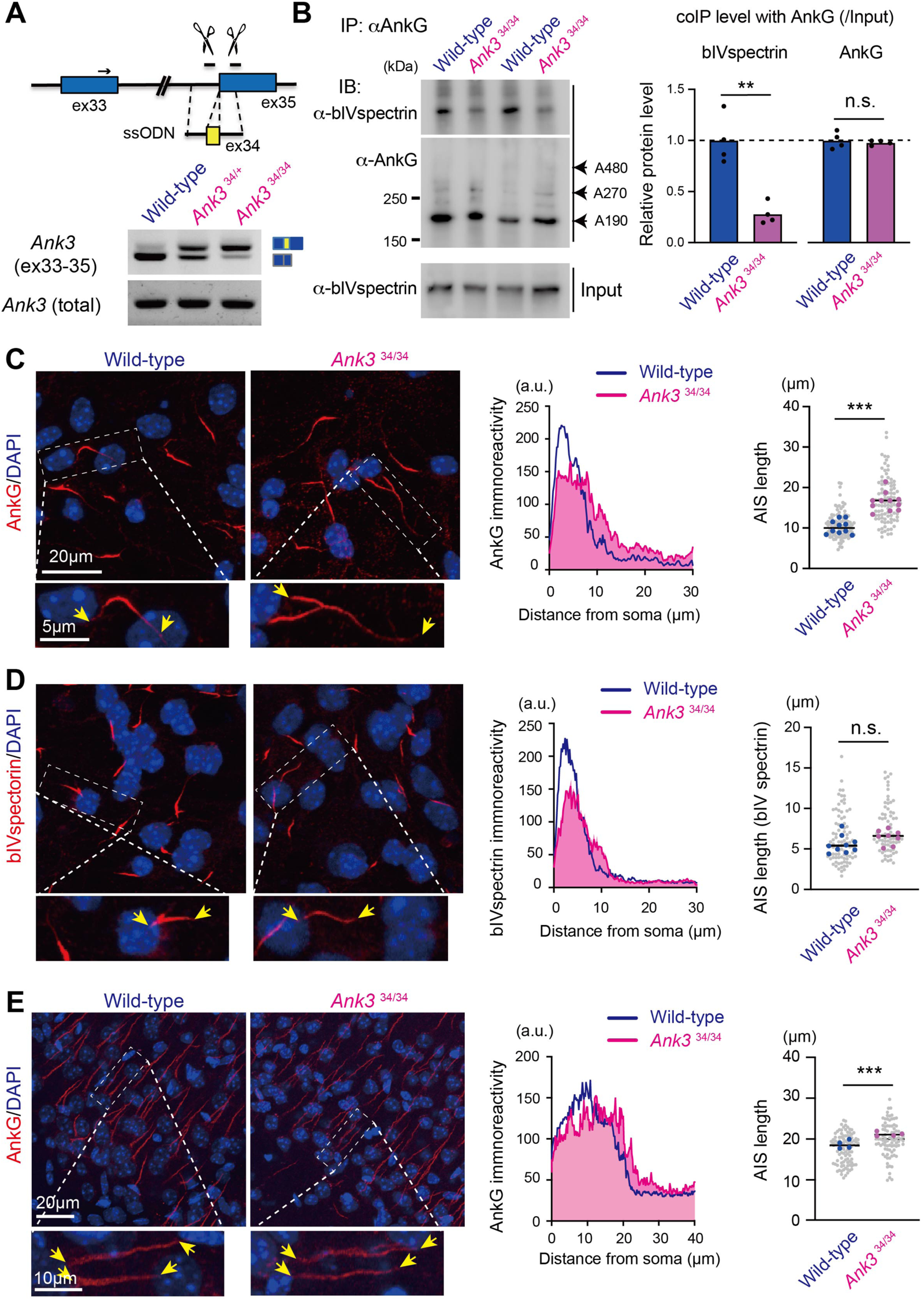
The constitutive insertion of *Ank3* micro-exon 34 impairs the interaction of AnkG and βIV spectrin and induces AIS structural changes in mice. (A) Schematic diagram demonstrating the constitutive insertion of micro-exon 34 at *Ank3* genome using CRISPR/Cas9-mediated genome engineering (upper). Semi-quantitative RT-PCR images confirming the insertion of exon 34 in the *Ank3* gene. (B) Immunoblot showing coimmunoprecipitation of βIV spectrin and AnkG in cerebellar lysates of wild-type and Ank3 ex34 KI mice (left) and densiometric quantification (right). Immunoprecipitation was performed with anti-AnkG antibodies and proteins in inputs and immunoprecipitants were detected with anti-βIV spectrin and anti-AnkG antibodies. (n = 4). (C and D) Immunostaining of AnkG (C) or βIV spectrin (D) in CGCs of *Ank3* exon 34 KI mice and controls (left). Representative fluorescence intensity (middle) and quantification of AIS length (right) in both genotypes. The intensities of more than 10 axons were averaged per each genotype. (C) AnkG: n = 10 and 11 fields for wild-type and Ank3 ex34 KI mice, respectively. (D) βIV spectrin. n = 9 and 8 fields for wild-type and Ank3 ex34 KI mice, respectively. Unpaired t-test. (E) Immunohistostaining of AnkG in the cerebral cortex (motor area) of *Ank3* exon 34 KI mice and controls (left). Representative fluorescence intensity (middle) and quantification of AIS length (right) in both genotypes. n = 4 animals per genotype. unpaired t-test. High magnification images of the area surrounded by dashed rectangles in (C-E) are shown and the arrowheads mark the start and the end of the AIS. The scale bars represent 20 μm in (C, D, low magnification and E), and 5 μm in (C and D, high magnification).

Moreover, we extended our investigation of AIS structure to other brain regions of *Ank3* ex34 KI mice. In layer 2/3 of the motor cortex of *Ank3* ex34 KI mice, we observed elongated distribution and reduced intensity of AnkG, resembling the changes observed in CGCs (Figure 6E). These findings suggest that the significant impact of the insertion of micro-exon 34 at AnkG on AIS structure alterations extends beyond cerebellar neurons and may be implicated in homeostatic plasticity in other brain regions.

## Discussion

Homeostatic plasticity serves as a critical mechanism for sustaining brain activity and the dynamic regulation of AIS represents a key form of plasticity in the brain. The present study identified a crucial role of activity-regulated AS of small/micro-exons in the structural plasticity of cerebellar AIS.

### Control of late-onset AIS plasticity by nuclear Rbfox1-dependent AS of small/micro-exons

In this study, we demonstrated the involvement of neuronal activity-dependent AS in the plasticity of AIS in CGCs. Although previous studies showed that several forms of AIS structural plasticity exhibit a slow and late onset regulation on the order of days (7), the underlying mechanism governing this process has remained largely elusive. We have shown that high K^+^-induced depolarization can induce structural and functional plasticity in CGCs with the order of days, which could serve as a new model to elucidate the underlying mechanism governing late-onset AIS plasticity. The comprehensive transcriptomic analysis revealed that depolarization exhibits a remarkable tendency to alter the splicing of small exons including micro-exons (Figure 3B and 3C). We demonstrated that the removal of the activity-regulated exon 19 from Rbfox1 resulted in its nuclear accumulation and enhanced its splicing activity of the transcripts encoding AIS-enriched proteins such as *Nfasc* and *Scn8a.* These molecular changes were associated with the diffusion of AnkG distribution and alteration in AIS plasticity (Figure 4I). In contrast, *Rbfox* triple KO neurons differentiated from ES showed impaired AIS assembly (18). In our experiments, *Rbfox1* single KO CGCs did not exhibit significant alteration in AIS formation, possibly owing to the functional redundancy of other Rbfox family members. Based on this, we propose the idea that, while the Rbfox family is essential for AIS formation, the splicing of *Rbfox1* exon 19 may be critical for AIS plasticity.

Notably, the depolarization-induced exclusion of exon 19 occurred rapidly (<6hrs), and triggered subsequent changes in the AS of its target exons at later stages (>24hrs) over a span of days (Figure 4H). Therefore, the sequential AS events orchestrated by nuclear Rbfox1 likely represent a major mechanism driving late-onset molecular alterations during AIS plasticity. We propose that the late-onset plasticity, at least as observed in our study, could be explained by the requirements of newly synthesized proteins and their subsequent molecular events.

The Rbfox family regulates the splicing of multiple transcripts encoding AIS-enriched proteins such as ion channels (Na^+^ and K^+^ channels), CAMs (neurofascin, NrCAM), and scaffolding proteins (AnkB, AnkG) (18, 21, 22). Among them, our study has specifically identified a subset of nuclear Rbfox1-targeted exons in a neural activity-dependent manner. Notably, the splicing shift of exons 26-29 in neurofascin could potentially serve as a critical mechanism for mediating activity-mediated structural AIS change. This is because neurofascin recruits extracellular matrix (ECM) molecules to the AIS (34), thus stabilizing AIS via a connection between the AIS submembrane cytoskeleton and the ECM. Genome-wide mapping of Rbfox splicing events has been conducted previously with HITS-CLIP, exon array, and RNA-seq (18, 21, 22, 35, 36) but the micro-resolution mapping of the “activity-regulated” AS program by nuclear Rbfox1 would be worthwhile to characterize the distinct roles of AS isoforms in the plasticity events.

### The strong impact of activity-dependent micro-exon splicing at AnkG on structural AIS plasticity

Given the drastic changes in the cerebellar AIS following high K^+^ stimulation, it is evident that these plasticity changes cannot be solely attributed to nuclear Rbfox-mediated splicing alterations, suggesting the involvement of multiple regulatory mechanisms. We identified the Rbfox1-independent mechanism that involves the activity-dependent insertion of micro-exon 34 at AnkG, which notably impacts structural changes during AIS plasticity in mature neurons (Figures 5 and 6). We demonstrated that the insertion of exon 34 strikingly altered the distribution of the neuronal isoform of AnkG, AnkG480 (Figure 5), mirroring the depolarization-induced diffusion of endogenous AnkG (Figure 1). Similarly, the CGCs of *Ank3* ex34 KI mice exhibited diffused and elongated AIS, along with weakened interaction between AnkG and βIV spectrin. This is conceivable because exon 34 encodes a micro-segment of SBD and its insertion negatively affected the interaction between βIV spectrin and AnkG in cell lines (18, 20). In addition, βIV spectrin stabilizes other AIS-enriched proteins through its interaction with AnkG rather than initial assembly (37). Given our results that the insertion of exon 34 in AnkG endogenously disrupts its interaction with βIV spectrin (Figure 6), we speculate that activity-inserted exon 34 at AnkG could exert structural change in AIS plasticity of CGCs by destabilizing the assembly of several AIS proteins scaffolded by AnkG such as Na_v_ and neurofascin.

While a previous study showed that AS of *Ank3* is regulated by the Rbfox family-mediated splicing program (18), our data suggest that the activity-regulated insertion of exon 34 at *Ank3* in mature CGCs is unlikely to be regulated by a Rbfox1-dependent mechanism. The underlying core regulatory factors responsible for this Rbfox1-independent activity-induced late-onset AIS plasticity remain elusive, warranting further exploration. Investigating RNA-binding proteins that preferentially regulate AS of micro-exons in the mammalian brain could provide valuable insights. Additionally, emerging evidence highlights the potential roles of chromatin remodeling and RNA N6-methyladenosine (m6A) modification in AS (38, 39). A recent study underscored the influence of histone modification in activity-induced late-onset AS of neurexin-1, implicating its involvement in memory consolidation (40). Therefore, it would be of great interest to explore the role of these mechanisms in governing late-onset splicing and AIS plasticity in future studies.

### AS-dependent structural and functional changes at the AIS in cerebellar neurons as a novel form of AIS plasticity in the CNS

In the last decade, numerous studies have reported various forms of homeostatic AIS plasticity, each showing distinct structural changes (6–9). Our study revealed that AIS plasticity in CGCs is functionally identical to those reported previously, albeit differ in some aspects. Unlike, the activity-induced AIS shortening from the distal ends observed in hippocampal GC neurons and the NM neurons of avian cochlear nucleus neurons of neurons (14–16), increased activity diffusely elongated AIS in CGCs to dampen neuronal excitability through the reduced density of Na_v_ channels (Figures 1 and 2). Additionally, while high K^+^ stimulation altered AIS location from the soma in hippocampal CA neurons (11, 12), the AIS location in CGCs remained unchanged. Thus, this study introduced a newly novel type of AIS plasticity in CGCs, highlighting the complexity and diversity of homeostatic AIS plasticity across distinct brain areas /neuronal cell types. It would be necessary to explore whether the AS-dependent plasticity at the AIS is a physiological mechanism that occurs in other neuron types beyond CGCs.

Our findings raise intriguing questions regarding the implication of AS-dependent plasticity at the AIS in brain disorders and injuries. It would be necessary to investigate the link between AIS plasticity and physiological functions or neurological diseases. Notably, several mutations and variants of key regulatory genes for AIS plasticity such as *Rbfox1* and *Ank3* have been implicated in various neuropsychiatric disorders including autism spectrum disorder, bipolar disorder, and schizophrenia (41, 42). Investigating the impact of these mutations on the splicing of the relevant micro-exons and AIS plasticity could provide valuable insights into the underlying pathogenesis of these disorders and pave the way for the development of pioneering therapeutic strategies.

## Materials and Methods

### Antibodies

For immunoblot and immunostaining analyses, the following commercially available antibodies were used: mouse anti-ankyrinG (N106/65, Neuromab, Davis, CA, USA), rabbit anti-ankyrinG (Rb-Af610, Frontier Institute, Hokkaido, Japan), rabbit anti-panNav (Rb-Af800, Frontier Institute), rabbit anti-neurofascin (ab31457, Abcam, Cambridge, MA, USA), mouse anti-Rbfox1 (1D10, Millipore, Burlington, MA, USA), rabbit anti-GAPDH (G9545, Sigma Aldrich, St. Louis, MO, USA), mouse anti-PSD-95 (K28/43, Neuromab), mouse anti-TuJ1 (Tubulin b3) (MMS-435P, Covance, Princeton, NJ, USA), mouse anti-MAP2 (AP20, Millipore), and rat Anti-GFP (04404-84, Nacalai Tesque, Kyoto, Japan). The rabbit anti-βIV spectrin was previously described (43).

### Neuronal cell culture, the pharmacological and knockdown experiments

(19, 35)The CGCs were prepared from postnatal ICR mice aged 5-7 days by tissue dissociation utilizing 0.05% trypsin (Sigma Aldrich, St. Louis, MO, USA) and DNase I (Roche Applied Science, Penzberg, Germany) at 37°C for 10 minutes. The trypsin activity was inactivated with a soybean trypsin inhibitor (Sigma Aldrich). The cells were then seeded onto poly-D-lysine-coated dishes at a density exceeding 2.0 × 105 cells/cm2 and cultured for 12-15 days in Neurobasal medium (Invitrogen, Waltham, MA, USA) supplemented with 2% B27, 2 mM Glutamax, and penicillin/streptomycin (Invitrogen). Pharmacological experiments were conducted using Bicuculine, Nimodipine, U1026, and STO609 (TOCRIS, Bristol, UK). Knockdown experiments involved the use of Accell non-targeting control siRNA (D-001910-01, GE Healthcare Dharmacon, Lafayette, CO, USA) and Accell mouse Rbfox1 siRNA (A-041929-13, GE Healthcare Dharmacon). The efficacy of knockdown was validated in previous studies (19, 35).

### Immunostaining for culture, image acquisition, and analysis

Cultured cerebellar neurons were fixed using 4% paraformaldehyde in ice-cold phosphate-buffered saline (PBS) for 15 minutes. Subsequently, the neurons were permeabilized with 0.15% TritonX-100 in PBS for 15 minutes at room temperature and then incubated in a blocking solution (5% normal goat serum in PBS) for a minimum of 30 minutes at room temperature. Primary antibodies were added and left to incubate for 24 hours at 4°C. For visualization, appropriate secondary antibodies conjugated to Alexa 546 or 488 (goat, 1:1000) (Life Technology) were employed. Neuron morphology was visualized by staining for a neuronal marker, βIII-tubulin (TuJ1), or MAP2, with nuclei highlighted by DAPI staining. For image acquisition and analysis, confocal imaging was conducted using the LSM700 confocal system (Zeiss, Oberkochen, Germany). ImageJ software (National Institutes of Health: Bethesda, MD, USA) was used for the analysis of the original images. After the appropriate threshold (top 2.5%– 5.0% of display range) was set as illustrated in Figure 1B, the length or intensity of AIS immunoreactivity was measured on the axons of CGCs. The length of AIS (approximately 4-23 axons) within 1 x 10^4^ um^2^ areas (1 field) was measured and averaged as a single point unless otherwise mentioned. The data was collected from three independent cultures or more.

All procedures related to the care and treatment of animals were strictly adhered to the Guide for the Care and Use of Laboratory Animals of Tokai University. The mice were housed under specific pathogen-free conditions at the Laboratory Animal Center, Tokai University. The experimental protocol was approved by the Institutional Animal Care and Use Committee of Tokai University (permit number 235022). all surgeries were performed under sodium pentobarbital anesthesia, with every effort made to minimize animal suffering.

### Immunoblot analysis

Cells or brain tissues were lysed with RIPA buffer (25 mM Tris-HCl, pH 8.0, 150 mM NaCl, 1% NP-40, 1% deoxycholate, 0.1% SDS) containing a protease inhibitor cocktail (Roche Applied Science, Penzberg, Germany). For protein interaction studies, the soluble fractions were subjected to immunoprecipitation for 24 h at 4°C and were subsequently examined via immunoblotting. A horseradish peroxidase-conjugated secondary antibody and enhanced chemiluminescence detection (Pierce, Dallas, TX, USA) were employed for visualization, with the signals captured using an image analyzer (LAS500; GE Healthcare).

### RNA isolation, alternative-splicing assays, and RT-qPCR

RNA was extracted using Nucleospin RNA XS (TaKaRa, Tokyo, Japan) or RNAiso Plus reagent (TaKaRa), followed by the removal of residual DNA with the Turbo DNA-free kit (Ambion, Austin, TX, USA). Two micrograms of the RNA were subjected to reverse transcription with random hexamers and the PrimeScript™ 1st strand cDNA Synthesis Kit (TaKaRa). For semi-quantitative PCR, DNA fragment intensities were quantified using an image analyzer (FAS-III, Toyobo, Osaka, Japan) and the ImageGauge software (Fujifilm, Valhalla, NY, USA). The intensity value for each splicing isoform band was standardized to the total transcript value. Similarly, reverse transcription-quantitative PCR (RT-qPCR) was conducted on a StepOnePlus qPCR system (Applied Biosystems, Waltham, MS, USA) using the Power SYBR Green PCR Master Mix (Applied Biosystems) and the comparative CT method. The transcript levels were normalized to the *Gapdh* reference gene. To prevent bias from varying total transcript amounts between groups, the transcript levels of each splicing isoform were also normalized to the total transcripts, as previously described. The oligonucleotide primer sequences used for both semi-quantitative PCR and RT-qPCR can be found in Tables S1 and S2.

### RNA-seq analysis

The RNA was extracted from CGCs treated with high K+ for 3 days from DIV9-DIV12 and untreated controls. After the QC procedures, mRNA from eukaryotic organisms was enriched using oligo(dT) beads and then rRNA was removed using the Ribo-Zero rRNA Removal Kit (Illumina, San Diego, CA, USA). First, the enriched mRNA was fragmented randomly using, fragmentation buffer, followed by cDNA synthesis using NEB Next Ultra RNA Library Kit (Illumina) according to the manufacturer’s protocols. The qualified libraries were pooled and fed into NovaSeq6000 sequencers (Illumina) according to the recommended pooling protocol based on effective concentration and anticipated data volume. The filtering process for cleaning reads included the removal of reads containing adapters, those with N > 10% (where N represents an undetermined base), and those with low-quality bases (Qscore <= 5) accounting for over 50% of the total base. The clean reads were aligned to the mouse reference genome mm10 using the hisat2 program (44). Differentially spliced AS exons were identified and quantified using vast-tools. The exon alteration was represented with delta PSI (dPSI), calculated as the difference between the average PSI of the high K+-treated group and the control group. The p-value was calculated using the PSI distribution of each group as input. The p-value for this event was found to be less than 0.0001, given the clear separation of the two distributions. The data presented in this manuscript have been deposited in NCBI’s Gene Expression Omnibus and are accessible through GEO Series accession number GSE246175.

### The construction and production of lentivirus vectors for shRNA and gRNAs

For the knockdown of Ankyrin-G, lentivirus encoding small hairpin RNA (shRNA) was prepared using the plentiCMV vector (RIKEN BioResource Research Center) with the following shRNA sequences: 5’-CTCAGTATGACTCAAGGTTTC-3’ for AnkG exon 34 (+); 5’-GCATTCTGGGTTTCTGGTTAG-3’ for AnkG exon 34 (-); 5’-CCTGCTCATAGGAAGAGGAAA-3’ for AnkG exon 1 (total); 5’-GCTGAGTACTTCGAAATGTC-3’ for control (Luciferase). For the design of the lentiviral vectors for the *in vitro* genome editing, the sequences of gRNAs were selected using CHOPCHOP (https://chopchop.cbu.uib.no) (>Rank5) (Table S3). These sequences were inserted into the U6 promoter-driven lentiviral vector using the Lentiviral CRISPR/Cas9 System (CASLV511PA-G System Biosciences, Palo Alto, CA, USA), following the manufacturer’s protocol. The lentiviral vectors were generated by transfecting human embryonic kidney cells (HEK293T) with a mixture of four plasmids using a calcium phosphate precipitation method. The four-plasmid mixture consisted of 6 μg of pCAG-kGP1R, 2 μg of pCAG-4RTR2, 2 μg of pCAG-VSV-G, and 10 μg of the vector plasmid Carrying the gRNA or the shRNA. After 40 hours, the medium containing virus particles was concentrated by centrifugation at 15,000g for 5 to 7 h.

The resulting virus particles were then suspended in cold PBS (pH 7.4), frozen in aliquots, and stored at -80°C until use. After assessing the titer, the appropriate amount of lentivirus was transduced into cultured neurons 5 to 7 days before harvesting the cells.

### Construction of helper-dependent adenovirus vector (HdAdV) expressing AnkG

The plasmid used for the production of a HdAdV was constructed to carry an inverted terminal repeat (ITR) sequence, adenovirus packaging sequence (Ψ), and about 10kb of the human hypoxanthine phosphoribosyltransferase (HPRT) gene for size adjustment, with ITR at both ends. To incorporate the Cre-dependent expression unit into the plasmid, a neomycin resistance gene flanked by loxP sequences was inserted downstream of the CAG promoter, followed by a cloning site and the rabbit beta-globin poly A signal (45). AnkG cDNAs, both with and without exon34, were inserted into the cloning site and excised with NotI outside both ITRs for HdAdV preparation. The helper virus, an E1 and E3 deficient adenovirus vector, included a reverse Ψ flanked by two FLP target sequence FRTs, preventing the packaging of the helper virus genome into the virus capsid (46). The plasmids for HdAdV preparation containing AnkG were transfected into FLP-expressing 293 cells using lipofectamine LTX. After 24 hours, the helper virus was infected, and 72 hours later, HdAdVs were generated.

### Generation of Ank3 ex34 Knock-in mice

The mice were generated using the CRISPR/Cas9 system. The guide RNAs (gRNAs) and single-stranded oligodeoxynucleotides (ssODN) were designed as described previously (18) and purchased from IDT (Coralville, IA, USA). The Cas9, gRNAs, and ssODN were mixed and delivered to pseudopregnant female mice at embryonic day 0.5-1.5 (E0.5-E1.5) using *i*-GONAD (improved genome editing via oviductal nucleic acids delivery) method (47). The successful generation of the mice was confirmed through the conventional genotyping method and Sanger sequencing. The F0 mice subsequently crossed with wild-type mice for more than five generations to minimize potential off-target effects. The sequence for gRNA and ssODN were listed in Table S3.

### Electrophysiology

Whole-cell patch-clamp recordings were conducted using a Multiclamp 700B (Molecular Devices, San Jose, CA, USA) at a recording temperature of 29°C–30°C. Artificial cerebrospinal fluid (in mM: 125 NaCl, 2.5 KCl, 26 NaHCO3, 1.25 NaH2PO4, 2 CaCl2, 1 MgCl2, 17 glucose, pH 7.3) was perfused in the bath. Excitatory and inhibitory synaptic currents were blocked using 20 µM 6,7-dinitroquinoxaline-2,3-dione (DNQX, Sigma) and 10 µM SR95531 (Abcam), respectively. Pipettes were pulled from glass capillaries (GC-150TF, Harvard Apparatus, Holliston, MA, USA) with a P-97 horizontal puller (Sutter), and had a resistance of 7–9 MΩ when filled with a K+-based internal solution (in mM: 113 K-gluconate, 14 Tris2-phosphocreatine, 4.5 MgCl2, 4 Na2-ATP, 0.3 Tris-GTP, 0.2 EGTA, and 9 HEPES-KOH, pH 7.2). Series resistance (15–30 MΩ) was left uncompensated, and the liquid junction potential (–12 mV) was not corrected. Signals were low-pass filtered at 10 kHz and digitized at 100 kHz using an NI PCIe-6251 data-acquisition board via BNC-2090A (National Instruments, Austin, TX, USA). Data were acquired and analyzed offline using Axograph

### Statistical analysis

GraphPad Prism 10 (GraphPad Software, Inc., La Jolla, CA, USA) was used for the majority of the statistical analyses. Pairwise comparisons were performed using a Student’s t-test. For multiple comparisons, an analysis of variance followed by Turkey’s or Dunnet’s tests was used. Data are represented as the mean ± standard error of the mean. Significance is indicated as follows: ***, p < 0.001; **, p < 0.01; *, p < 0.05.

## Acknowledgments

We would like to thank Dr. Matthew Rasband (Baylor College of Medicine) for providing the βIV spectrin antibody and Dr. Takeshi Yoshimura (Osaka Univ.) for his insightful feedback on the manuscript. We would like to acknowledge all the members of the Support Center for Medical Research and Education at Tokai University for their assistance in conducting the experiments and maintenance of experimental animals. The work was supported by JSPS KAKENHI (grants 15H04277, 15K14355, and 20H03344), the Takeda Science Foundation, the Naito Foundation, the Novartis Foundation for the Promotion of Science, and Mochida Memorial Foundation for Medical and Pharmacological Research (all to T.I.).

