## Supplementary file for "Activity-regulated micro-exon splicing programs underlie late-onset plasticity at the axon initial segment"

**Supplementary Figures**

**Figure S1: Activity-dependent structural changes at the AIS in CGCs.**

Additional information related to Figure 1.

**Figure S2: RNA-seq analysis at the gene level in high K^+^-treated CGCs.**

Additional information related to Figure 3.

**Figure S3: RNA-seq analysis of alternatively spliced exons of genes encoding AIS proteins in high K^+^-treated CGCs.**

Additional information related to Figure 3.

**Figure S4: Activity-dependent *Rbfox1* exon19 skipping regulates the splicing of genes encoding AIS-enriched proteins.**

Additional information related to Figure 4.

**Figure S5: Mismatched axonal localization of neurofascin and βIV spectrin in the CGCs of *Ank3* ex34 KI mice.**

Additional information related to Figure 6.

**Supplementary Tables**

**Table S1. Oligonucleotide sequences for semi quantitative RT-PCR primer sets**

**Table S2. Oligonucleotide sequences for quantitative RT-PCR primer sets**

**Table S3. Oligonucleotide sequences for gRNAs and ssODN in CRISPR/Cas9 system**

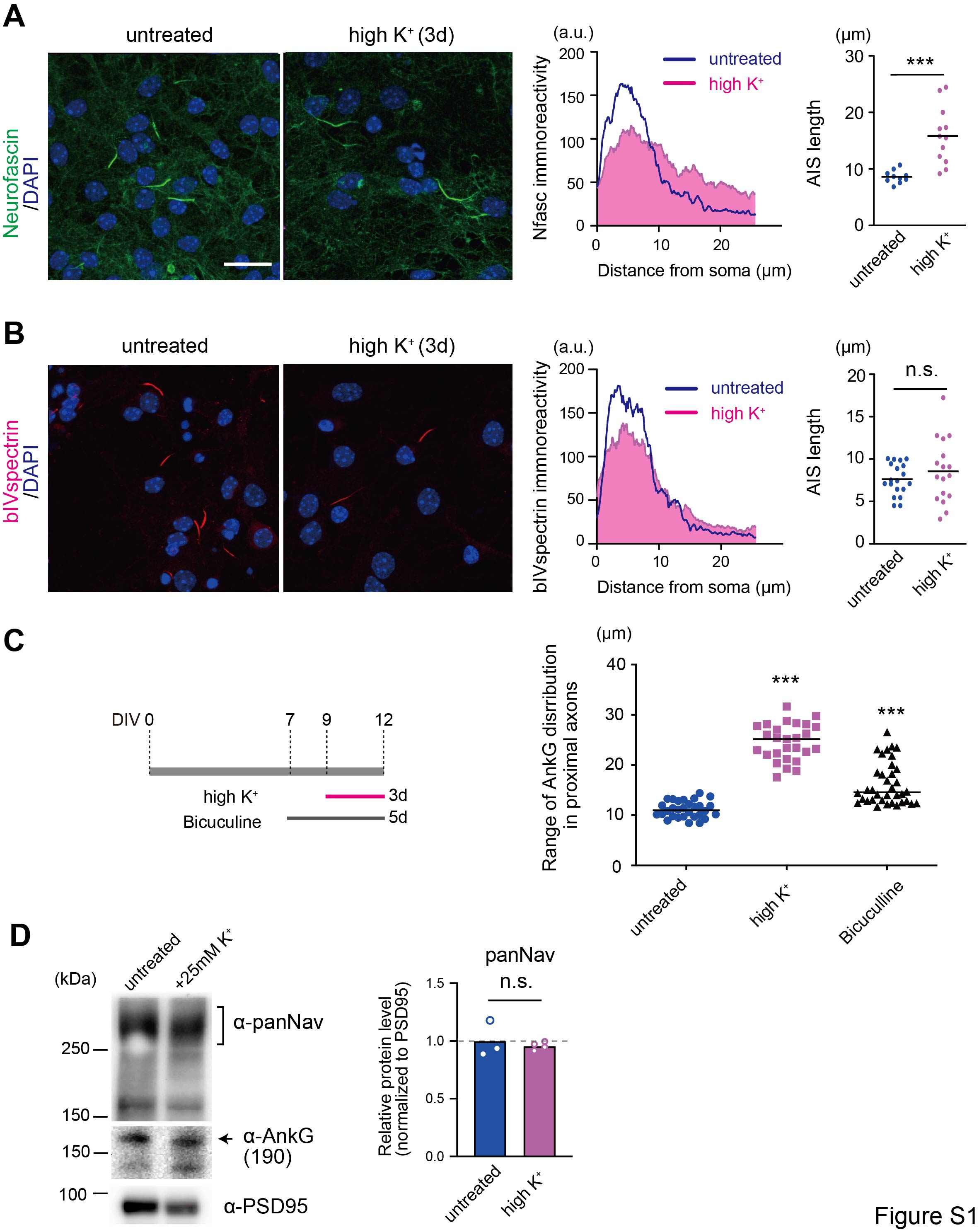

**Figure S1: Activity-dependent structural changes at the AIS in CGCs**

(A-B) Immunostaining of Neurofascin (A) and βIV spectrin (B) in untreated and high K+-stimulated CGCs for 3 days (3d) A. DAPI stains the nucleus (left). Representative fluorescence intensity (middle) and quantification of AIS length (right). The intensities of more than 10 axons were averaged per group. [n = 10-19 fields per group. unpaired t-test. (C) Schematic representation showing the time course of the *in vitro* pharmacological experiment (left). Quantification of AIS length (right). n= 30, 27 and 36 fields for untreated, high K^+^ and bicuculine, respectively. One-way ANOVA with Tukey’s multiple comparisons test. (E) Western blot showing the detection of ankG and panNav in untreated and high K+-stimulated CGCs. Densiometric quantification of panNav (right). n = 4 cultures. Unpaired t-test. The scale bars represent 20 μm in (A and B)

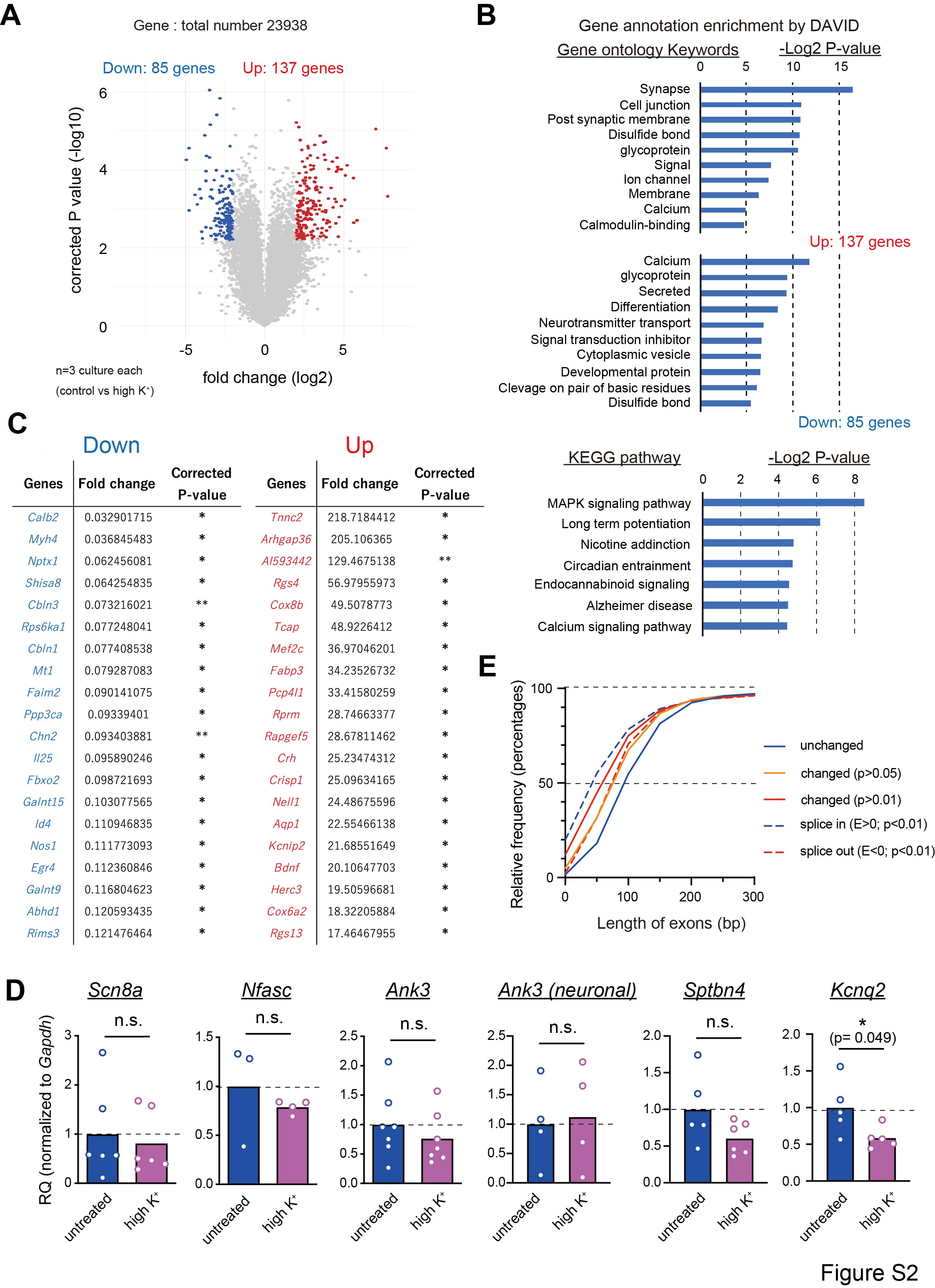

**Figure S2: RNA-seq analysis at the gene level in high K^+^-treated CGCs**

(A) Volcano plot showing the differentially expressed genes (DEGs) (high K^+^ vs control; threshold set: FC>4.0, corrected *p* value < 0.05). (B) Gene ontology (GO) analysis of the DEGs with high K^+^ stimulation by DAVID (Database for Annotation, Visualization and Integrated Discovery) functional research annotation (<https://david.ncifcrf.gov>). Enrichment was thresholded by corrected P-value (*p* <0.05). X-axis is represented as -log2 (P-value). The top terms of GO (upper) and KEGG-pathway (lower) are listed. (C) Lists of top upregulated and downregulated DEGs with FC. (D) Semi-quantitative RT-PCR quantification of genes encoding AIS-enriched proteins. n = 4-6 cultures. Unpaired t-test. (E) The cumulative distribution curve showing the relative frequency of exon length in unchanged and significantly altered exons with high K-treatment CGCs. The distribution curve of significantly altered exons shifted toward the left.

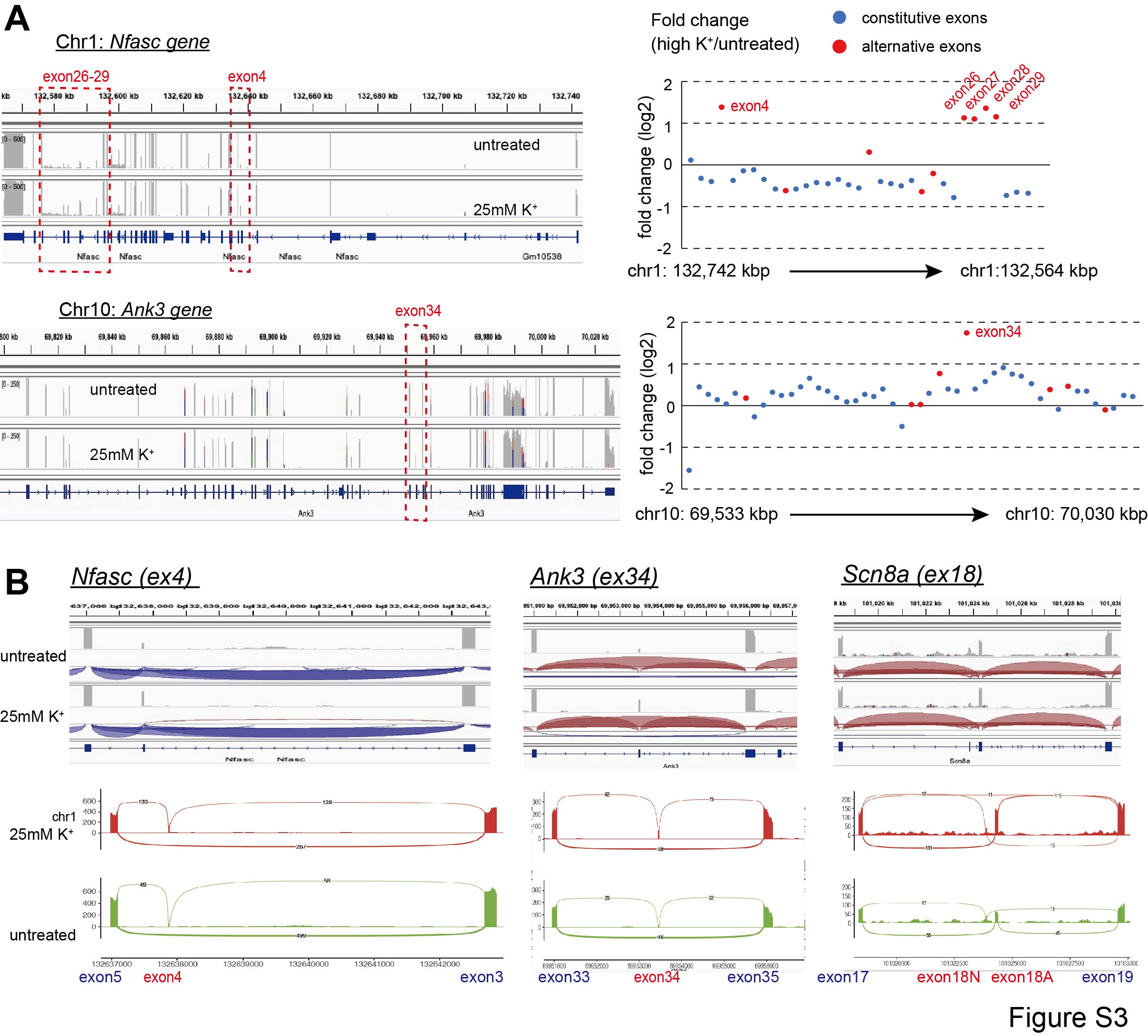

**Figure S3: RNA-seq analysis of alternatively spliced exons of genes encoding AIS proteins in high K^+^-treated CGCs**

(A) Specific alterations in alternatively spliced exons at *Nfasc* and *Ank3* genes. RNA-seq data of each gene set was analyzed by JunctionSeq software (<http://hartleys.github.io/JunctionSeq>) and modified. Blue and red dots represent the alternative and constitutive exons, respectively).

(B) Representative RNA-seq data and Sashimi plots (lower) of the alternatively spliced exons in *Nfasc*, *Ank3* and *Scn8a*. Each plot includes cassette exons and intron retentions. The *red plots* high K-treated CGCs and the *green plots* represent untreated CGCs. The *X*-axes show genomic loci, and the *Y*-axes indicate transcription intensity. A “sashimi-like” region in each plot indicates an exon region, and the blank regions between them indicate intronic regions. The numbers on the bridges crossing exons indicate junction reads. (<https://github.com/guigolab/ggsashimi>)

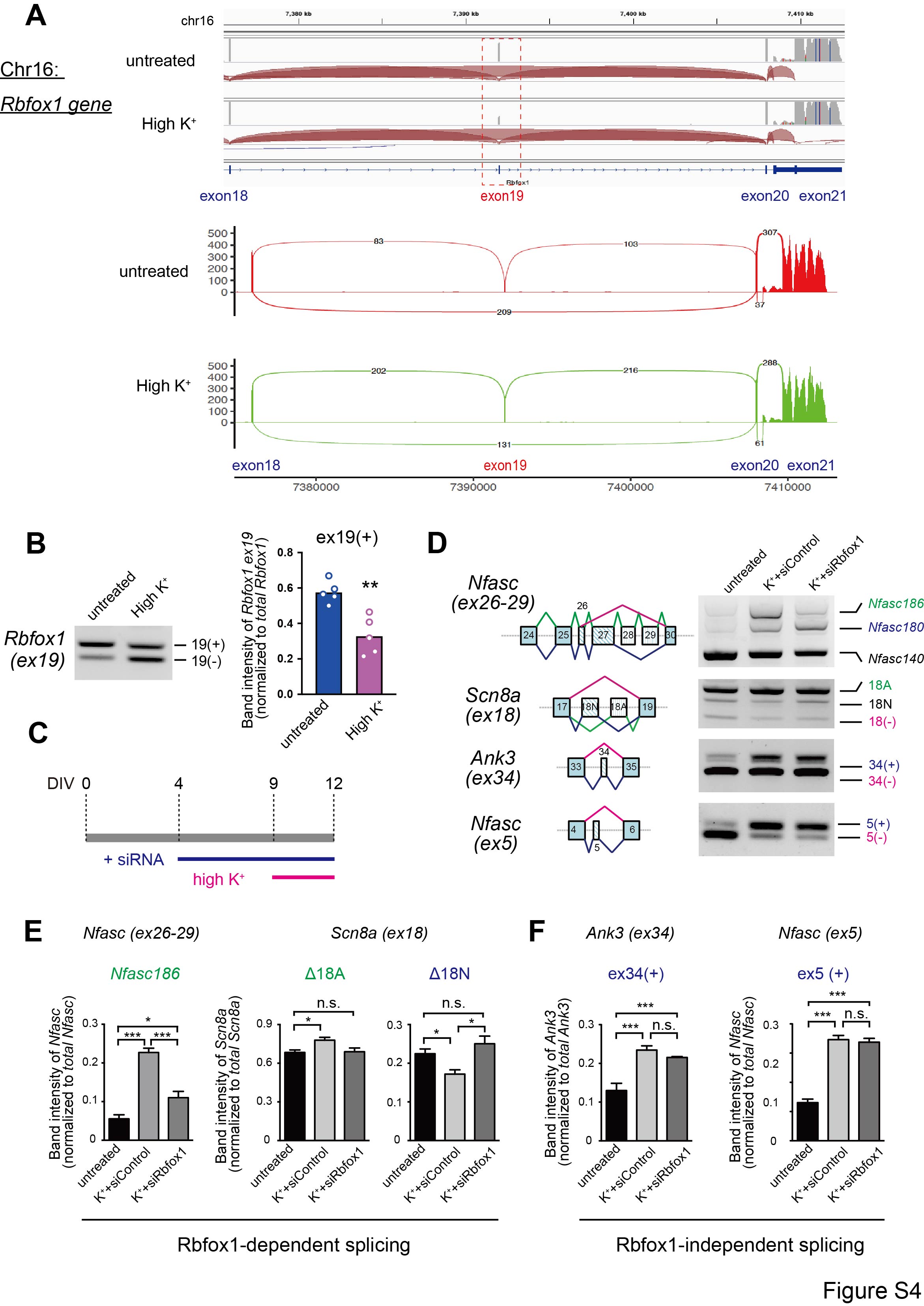

**Figure S4: Activity-dependent *Rbfox1* exon19 skipping regulates the splicing of genes encoding AIS-enriched proteins.**

(A) RNA-seq data and sashimi-plot of alternative exon 19 of *Rbfox1* genes foloowing high K^+^ treatment. (B) Semi-quantitative RT-PCR images (left) and densitometric quantification (right) of Rbfox1 exon 19 in untreated and high K-treated CGCs. n = 5 cultures per group, unpaired t-test. (C) Schematic representation showing the time course of the experiment. The CGCs were stimulated with high K^+^ (red bar) or treated with 1ug of cell-permeable small interfering ribonucleic acids (siRNAs) for Rbfox1 or scramble (control). (D) Semi-quantitative PCR images demonstrating the splicing changes in *Nfasc*, *Scn8a*, and *Ank3* in CGCs, either untreated or treated with high K+ stimulation with Rbfox1 siRNA or scramble siRNA. (E and F) Densiometric quantification of Rbfox1-dependent (E) and Rbfox1-independent spliced exons (F). n = 4-5 cultures. One-way ANOVA with Tukey’s multiple comparisons test.

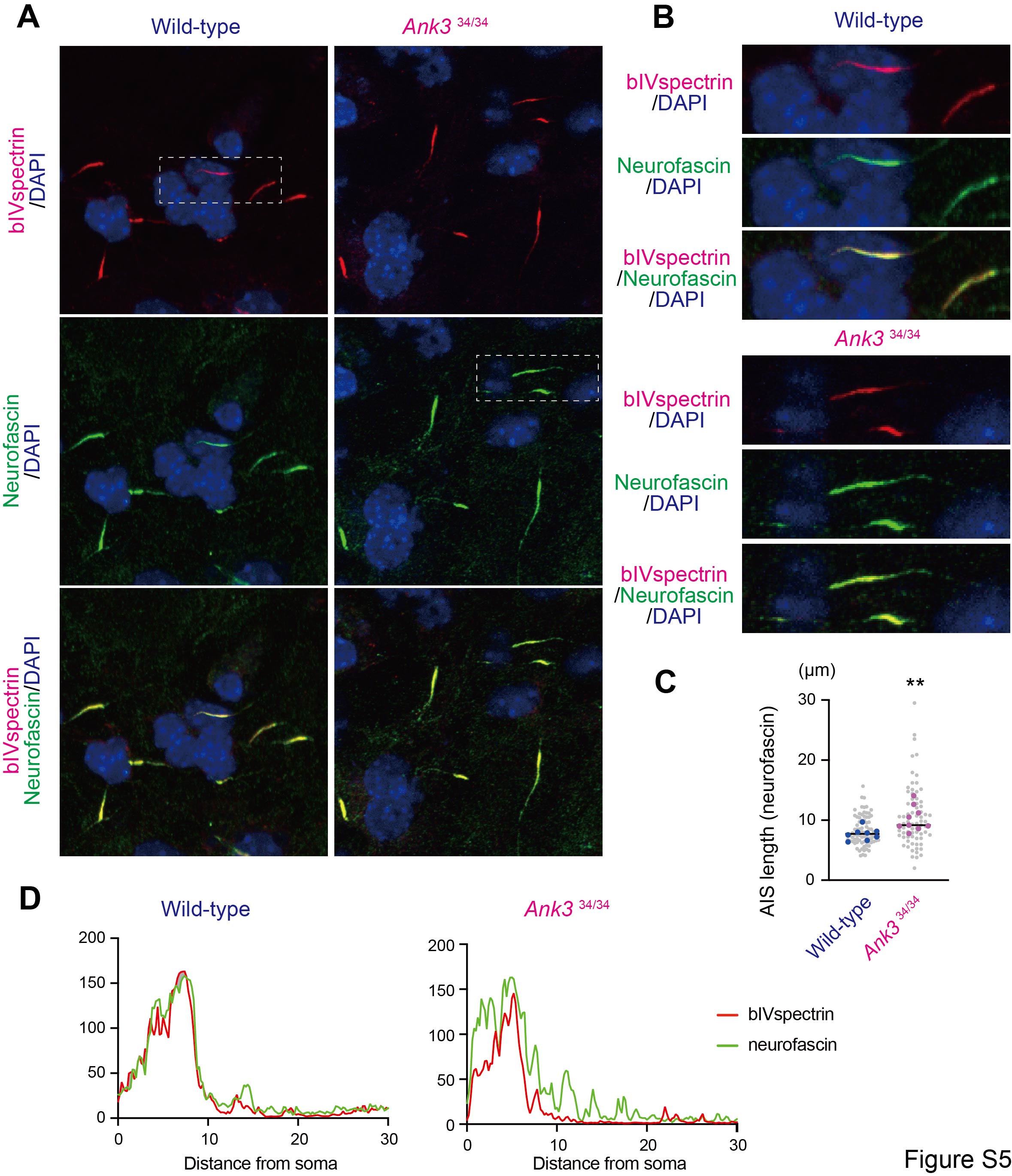

**Figure S5: Mismatched axonal localization of neurofascin and βIV spectrin in the CGCs of *Ank3* ex34 KI mice.**

(A) Co-Immunostaining of neurofascin and βIV spectrin in CGCs of *Ank3* exon 34 KI mice and controls. (B) High magnification images of the area surrounded by dashed rectangles in (A). (C) Quantification of AIS length represented by neurofascin immunoreactivity in both genotypes. n = 9 fields per each genotype, Unpaired t-test.
(D) Representative fluorescence intensity of neurofascin and βIV spectrin overlay in *Ank3* exon 34 KI mice and controls. The scale bars represent 10 μm in (A and B).

**Table S1: Oligonucleotide sequences for semi quantitative RT-PCR primer sets**

| Primer  (Forward)  (Reverse) | Sequence (5'-3') |
| --- | --- |
| Nfasc ex24-F  Nfasc ex30-R | 5'- ATA CAT CCT CAG ATA CGT GC -3'  5'- TGC CTG GTT ATT GGT GTA AG -3' |
| Scn8a ex7-F  Scn8a ex9-R | 5'- GCC TAT GGC TTC GTC AAG TT-3'  5'- ATG AGA CAC ACC AGC AGC AC -3' |
| Kcnq2 ex10-F  Kcnq2 ex12-R | 5'- CAC AGC CAG AGC CAT CAC CAA -3'  5'- CTC GGG CTG TCA TCA AGA CTC TGA TC -3' |
| Ank2 ex34-F  Ank2 ex34-R | 5'- CAC AGC CAG AGC CAT CAC CAA -3'  5'- CTC GGG CTG TCA TCA AGA CTC TGA TC -3' |
| Ank3 ex33-F  Ank3 ex35-R | 5'- GTT GGG CAG CGG ACA CGT TAG -3'  5'- CCT CGC ATG GAG CCC CCT C -3' |
| Nfasc ex3-F  Nfasc ex5-R | 5'- TCC ACA TAG CCC TCA TCC TC -3'  5'- TCA CGG ACT GCT TGG TGA TA -3' |
| Ndel1 ex8-F  Ndel1 ex10-R | 5'- TGT GGG GTA ATG AAC AGC AA -3'  5'- CAC ACA CTG AGA GGC AGC AT -3' |
| Rbfox1 ex17-F  Rbfox1 ex19-R | 5'- GTG GTT ATG CTG CGT ACC G -3'  5'- GGA GCA AGT GTG TGG TGG TA -3' |
| GAPDH-F  GAPDH-R | 5'- TGT TGC CAT CAA TGA CC -3'  5'- TCT CAT GGT TCA CAC CCA -3' |

**Table S2: Oligonucleotide sequences for quantitative RT-PCR primer sets**

| Primer  Forward (F)  Reverse (R) | Sequence (5'-3') |
| --- | --- |
| Scn8a ex17-F  Scn8a ex18N-R  Scn8a (total)-F  Scn8a (total)-R | 5'- GCC TAT GGC TTC GTC AAG TT -3'  5'- ACA CCC ACA GGT TCT TCA GC -3'  5'- CTG AAC AGC ACA GCC ATC AG -3'  5'- ACA CCC ACA GGT TCT TCA GC -3' |
| Nfasc (total)-F  Nfasc (total)-R  Nfasc ex27-F  Nfasc ex28-R | 5'- ACC TGG AGA CCA TCA ACC TG -3'  5'- TTT TCC ACC ATC TGC TTT CC -3'  5'- AGA GCC CTC CCA CTA CCA CT -3'  5'- GGT GAT GTT GGC CCA TTT AC -3' |
| Ank3 ex34- F  Ank3 ex35- R  Ank3 (total)-F  Ank3 (total)-R | 5'- GTC TCC GCT GCC TCA GTA TG -3'  5'- CAT ACT GAG GCA GCG GAG AC -3  5'- ACC TGG AGA CCA TCA ACC TG -3'  5'- TTT TCC ACC ATC TGC TTT CC -3' |
| Kcnq2 (total)-F  Kcnq2 (total)-R | 5'- GGA TAG CGA CAC GTC CAT CT -3'  5'- CAT CCA GGT TCT CCT TGG AC -3' |
| Sptbn4 (total)-F  Sptbn4 (total)-R | 5'- GCG AGG TGG CTA GTG ACT ACA -3'  5'- CCG TTC ATC TCC TCC TCA TC -3' |
| GAPDH-F  GAPDH-R | 5'- TGT TCC AGT ATG ACT CCA CTC ACG -3'  5'- AGT AGA CTC CAC GAC ATA CTC AGC -3' |

**Table S3: Oligonucleotide sequences for gRNAs in CRISPR/Cas9 system**

| Primer  sgRNA(sg)1  sgRNA(sg)2 | Sequence (5'-3') |
| --- | --- |
| Rbfox1 ex19 -1  Rbfox1 ex19 -2 | 5'- TTC ACA ATA CTG CTC GCT CA-3'  5'- GGA ATG TCT GTT ACC TGC AG -3' |
| Ank3 ex35 -1  Ank3 ex35 -2 | 5'- TCC AAC CCC GCA CAG GTT TC -3'  5'- AGC TAA CCA GAA ACC TGT GC -3' |
| Ank3  exon34/35 ssODN | 5'-GTGACGCCCTCTGATCTTTTTCCAACCCCGCACAGGTGCTCGTCTCCGCTGCCTCAGTATGACTCAAGGTTTCTCGTTAGCTTTATGGTGGACGCGAGAGGGG  -3' (exon34) |
| Rbfox1 KO -1  Rbfox1 KO -2 | 5'- TTA CAC CAA ACA TTT GTC GG -3'  5'- CTG GGG CTG GCC GTC GGT CG -3' |
